# In pursuit of saccade awareness: Limited control and minimal conscious access to catch-up saccades during smooth pursuit eye movements

**DOI:** 10.1101/2025.07.01.662353

**Authors:** Jan-Nikolas Klanke, Sven Ohl, Almila Naz Esen, Martin Rolfs

**Affiliations:** Berlin School of Mind and Brain, Humboldt-Universität zu Berlin; Department of Psychology, Humboldt-Universität zu Berlin; Department of Psychology, Freie Universität zu Berlin; Bernstein Center for Computational Neuroscience Berlin, Humboldt-Universität zu Berlin

**Keywords:** Catch-up saccade, Pursuit eye movement, Motor control, Sensorimotor awareness

## Abstract

Observers use smooth pursuit to track moving objects—like koi carp gliding through a pond. When positional errors accumulate, rapid catch-up saccades correct for them. Despite their abruptness, these saccades usually go unnoticed, creating the seamless experience of smooth tracking. We conducted three experiments to examine awareness and control of catch-up saccades (**Experiment 1**), the effect of training (**Experiment 2**), and of movement intention (**Experiment 3**). All experiments followed a similar protocol. On each trial, a target moved horizontally at one of three constant speeds (3–12 dva/s). Two horizontal stimulus bands with vertically oriented gratings appeared above and below the trajectory. These bands were rendered invisible during pursuit by rapid phase shifts (>60 Hz), but became visible when briefly stabilized on the retina—either by a catch-up saccade or its replayed retinal consequence—providing immediate, saccade-contingent visual feedback. Observers reported whether they had seen the stimulus bands (visual sensitivity) and whether they were aware of making a catch-up saccade (saccade sensitivity). Visual sensitivity was consistently higher in trials with a catch-up saccade, confirming that these movements reduce retinal motion and enhance visibility. Higher target speeds increased saccade rate, but observers struggled to control them consciously: Visual feedback and training had no effect on the ability to control catch-up saccades. Only suppression-instructions yielded a small reduction. Saccade sensitivity was near zero, even in trials with saccade-contingent feedback. Neither training nor intention improved awareness. Together, our data suggest a limited ability to control and a low level of sensorimotor awareness of catch-up saccades during pursuit.

**Significance statement:** Smooth pursuit eye movements allow us to track moving objects seamlessly, yet these movements are frequently interrupted by small, rapid corrective catch-up saccades. Despite their disruptive nature, observers rarely notice their own catch-up saccades. To address this conundrum, we used a novel paradigm employing a stimulus that remains invisible during pursuit but becomes visible when retinally stabilized by a catch-up saccade—providing saccade-contingent visual feedback to investigate conscious control and awareness of catch-up saccades during pursuit. Our data show that higher target speeds increased saccade rates, and observers were largely unable to modulate this rate—except for a slight reduction when explicitly instructed to pursue as smoothly as possible. Moreover, awareness remained low even with saccade-contingent feedback, and neither training nor intention improved it. Together, these findings suggest limited conscious control and low sensorimotor awareness of catch-up saccades during smooth pursuit.

## Introduction

Picture a koi pond, where vibrant carp glide effortlessly beneath the surface. One koi, with a particularly striking pattern, catches your attention, and you begin to track its path through the shifting background of other colorful fish. To focus on the koi, and follow its motion through the water, your eyes engage in a behavior called ‘pursuit’—a slow, smooth rotation of the eyes that fixes your center of gaze on a moving target without loss in visual sensitivity (Schütz et al., 2008, 2009).* Consequently, pursuit perfectly explains your stable, and detailed (high-resolution) impression of the fish on its trajectory through the pond. However, initiating and maintaining pursuit is limited by reaction time, and target speeds are neither reached instantly nor sustained perfectly (Goettker & Gegenfurtner, 2021), leading to deviations between the intended and actual gaze positions. To correct for these position errors, observers frequently initiate catch-up saccades: rapid eye movements that realign the center of gaze with the moving target (De Brouwer et al., 2002). Although catch-up saccades are necessary for successful tracking, they clash with how we experience pursuit: when tracking the koi in its pond, our impression is not of a jerky or unstable fish, but of one that remains fixed at the center of gaze while gliding smoothly through the water. Likewise, we feel as though our eyes move continuously and smoothly with the fish, rather than being frequently interrupted by abrupt, ballistic shifts in gaze position. In this study, we examine the discrepancy between the objective presence of catch-up saccades in gaze behavior and the subjective experience of smooth tracking. We examine conscious control and sensorimotor awareness of catch-up saccades during pursuit.

When studying conscious eye movement control and awareness, catch-up saccades are particularly relevant edge-cases: Observers can consciously perform regular saccades with high temporal (Kinder et al., 2008; Wong & Shelhamer, 2012) and spatial (Kowler & Blaser, 1995) accuracy. Catch-up saccades are generated during pursuit, however, and therefore primarily driven by visual motion—much like pursuit itself (Krauzlis, 2004; Rashbass, 1961). Despite evidence that catch-up saccades are an automatic response to position (or velocity) errors during pursuit (De Brouwer et al., 2002; Nachmani et al., 2020), it remains unclear whether they are entirely beyond voluntary control or if some level of control can still be exerted. Our first research objective was, therefore, to test whether observers can suppress (or at least postpone) catch-up saccades, or if these movements are invariantly triggered when the conditions for their generation are met. Catch-up saccades are equally interesting when it comes to sensorimotor awareness: They are mostly reflexive eye movements that are frequent, small, and fast, and, hence, have the potential to escape awareness (much like spontaneous microsaccades, cf. Klanke et al., 2025). As movements that are accompanied by visual transients as well as clear markers of success (i.e., the shift in the tracked object’s retinal position from peripheral to foveal), however, they might also be generated with a heightened degree of conscious oversight. This ambivalence makes them ideal for exploring our second research objective: understanding observers’ awareness of their catch-up saccades and the factors that modulate sensorimotor awareness.

Here, we present the results of three experiments investigating control and awareness of catch-up saccades. In all experiments, we use a similar paradigm that required participants to pursue a moving target with their eyes. To minimize initial catch-up saccades, each trial began with 500 ms of fixation while the target was already in motion (Rashbass, 1961).

Participants followed the target once it crossed the fixation point, continuing for 1000 ms (**Fig. 1a and c**). To compare performance across speeds and capture a range of motion dynamics, the target moved at a single constant speed per block: 6, 9, or 12 dva/s in **Experiment 1**, and 3, 6, or 9 dva/s in **Experiments 2** and **3**. At the end of each trial, participants were asked whether they believed they had made a catch-up saccade, allowing us to assess each observer’s sensitivity to their own saccades.

**Fig 1.**
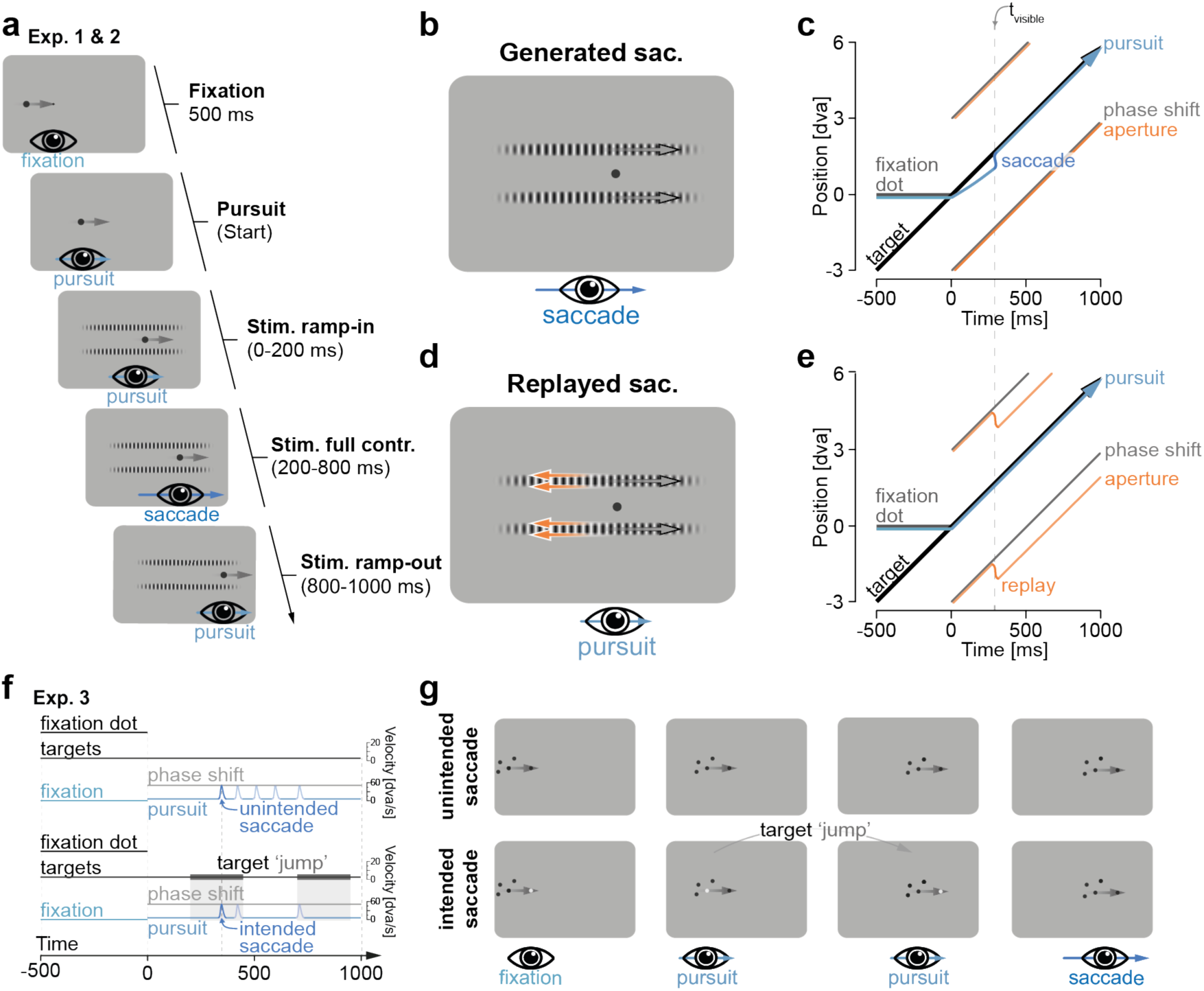
Experimental protocol and stimulus design. **a** Procedure of **Exp. 1** and **2**. Participants had to fixate for 500 ms while the movement target moved towards the fixation location at a constant speed of either 3, 6, 9, or 12 dva/s. They had to start pursuing, once the target fully occluded the fixation dot. Pursuit and stimulus interval lasted for 1000 ms, with stimulus bands increasing to 50 % contrast during the first 200 ms—and decreasing to 0 % in the 200 ms of each trial. **b** Stimulus display for generated catch-up saccades. Gray arrows indicate the direction of the phase shift; blue arrow indicates the direction of a catch- up saccade that leads to a retinal stabilization of the stimulus. **c** Spatiotemporal configuration of fixation dot, movement target, eye position, and stimulus (aperture and phase) during a trial (schematic, for actual gaze traces, velocity, and acceleration data see **Fig. M1**). The time of the saccade (tvisible) marks the moment when the generated saccade stabilizes the phase shift on the retina. **d** Stimulus display for replayed catch-up saccades. Gray arrows indicate the direction of the phase shift (as in panel b), while orange arrows indicate the direction of an aperture shift that replicates the retinal consequences of a saccade, resulting in retinal stabilization of the stimulus similar to the saccade shown in panel b. **e** Spatiotemporal configuration of fixation dot, movement target, eye position, and stimulus (aperture and phase) during a replay trial. The time of stimulus visibility (tvisible) is aligned with an aperture motion that replicates the retinal consequences of a saccade. **f** Procedure during unintended and intended catch-up saccade conditions in **Exp. 3**. In the unintended saccade condition, the procedure closely matched that of the first two experiments (e.g., without a target jump), with all saccades occurring unintentionally. In the intended condition, gray bands indicate the timing of the early or late target jumps (e.g., the brief recoloring of the current and future pursuit targets; see panel g for details). **g** Instruction conditions in **Exp. 3**. Unintended saccade condition (upper row): Participants were instructed to fixate on a black dot (panel 1) until one dot from the moving cloud crossed the fixation point (panel 2). They then tracked the black dot as it moved (panel 3), allowing spontaneous catch-up saccades to occur without explicit instruction (panel 4). Intended saccade condition (lower row): Participants were instructed to fixate on a white dot (panel 1) until it was crossed by a moving dot from the cloud, which turned white upon fully occluding the fixation dot (panel 2). They then tracked the moving white dot until it turned black, and another dot flashed white for 50 ms (panel 3), serving as the go-signal to make a catch-up saccade. Subsequent saccades were thus labeled intended.

To understand sensorimotor awareness of catch-up saccades, it is essential to consider the role of visual perception. When tracking a koi in a pond, we have the impression of continuously foveating the fish, without any noticeable disruptions. This perceptual continuity suggests that visual mechanisms contribute to masking the abrupt retinal shifts caused by catch-up saccades, preserving the illusion of smooth and uninterrupted pursuit. To assess the role of perceptual consequences in this process, we carefully controlled the amount of visual information available during catch-up saccades: We added a high-speed stimulus to the design that consisted of two vertical bands, placed 3 dva above and below the target trajectory. Each band had a high spatial frequency (5 cycles per degree; cpd) and a rapid phase shift (>56.50 dva/s), making it invisible during pursuit (**Fig. 1b**). However, a catch-up saccade in the same direction and with comparable peak velocity could stabilize the stimulus on the retina for a brief moment, providing immediate saccade-contingent feedback that informed the participant of their eye movement. To distinguish the perceptual consequences of eye movements from the motor effects, we introduced a replay condition, that used the same stimulus but added aperture motion, shifting the stimulus position similarly to a saccade, and creating a similar visual impression in the absence of a catch-up saccade (**Fig. 1e**). A no-stimulus condition served as an additional control for baseline perceptual reports and to ensure that any observed effects were due to the presence of the stimulus and not general task demands or expectations. To assess whether the stimulus provided saccade-contingent feedback as intended, participants were asked at the end of each trial whether they had perceived the stimulus and, if so, which one they had noticed (except in the first experiment). Their responses were used to calculate visual sensitivity.

To systematically investigate control and awareness of catch-up saccades, we conducted three experiments, each introducing a specific variation to isolate different contributing factors: **Experiment 1** was conducted to establish the paradigm and assess baseline sensitivity (both visual and saccadic). In **Experiment 2**, participants were instructed to suppress their catch-up saccades to determine whether they could be trained to voluntarily control these movements or learn to become more aware of them. We also recruited two groups: naïve observers with no prior eye movement study experience and experts with extensive knowledge of eye movements that had previously participated in multiple studies. This distinction was made to assess whether training effects were general or required a longer time to manifest (see **Supplementary Material** *S2: Observer groups in **Experiment 2*** for a detailed analysis). Additionally, we enhanced the saccade-contingency of the stimulus and informed participants that perceiving the stimulus likely resulted from a catch-up saccade, which they should use as immediate sensory feedback. In **Experiment 3**, we manipulated movement intention by presenting specific movement instructions in each trial. In pursuit trials, participants had to pursue the target without generating a catch-up saccade (**Fig. 1f and g**, upper row), while in saccade trials, participants were instructed to generate a specific saccade (**Fig. 1f and g**, lower row). Trial type was indicated at the start by the color of the fixation dot: white for saccade trials and black for pursuit trials. Both instructions were presented randomly interleaved within each block.

## Results

### Visual sensitivity to intra-saccadic stimulation

To assess whether the stimulus was visible in the absence of saccades, we calculated visual sensitivity in pursuit-only trials. In **Experiment 1**, sensitivity was significantly above zero (d’ = 0.47 ± 0.28, *p* = 0.005; **Fig. 2a**), suggesting that the high phase-shift velocity may have introduced speed-related aliasing artifacts that made the stimulus faintly visible during smooth pursuit. In contrast, sensitivity did not differ from zero in **Experiments 2** and **3** (**Exp**. **2**: d’ = 0.08 ± 0.25, *p* > 0.250; **Exp. 3**: d’ = 0.09 ± 0.17, *p* > 0.250; **Fig. 2a**), indicating that lowering the phase shift velocity effectively rendered the stimulus invisible in trials without catch-up saccades.

**Fig. 2.**
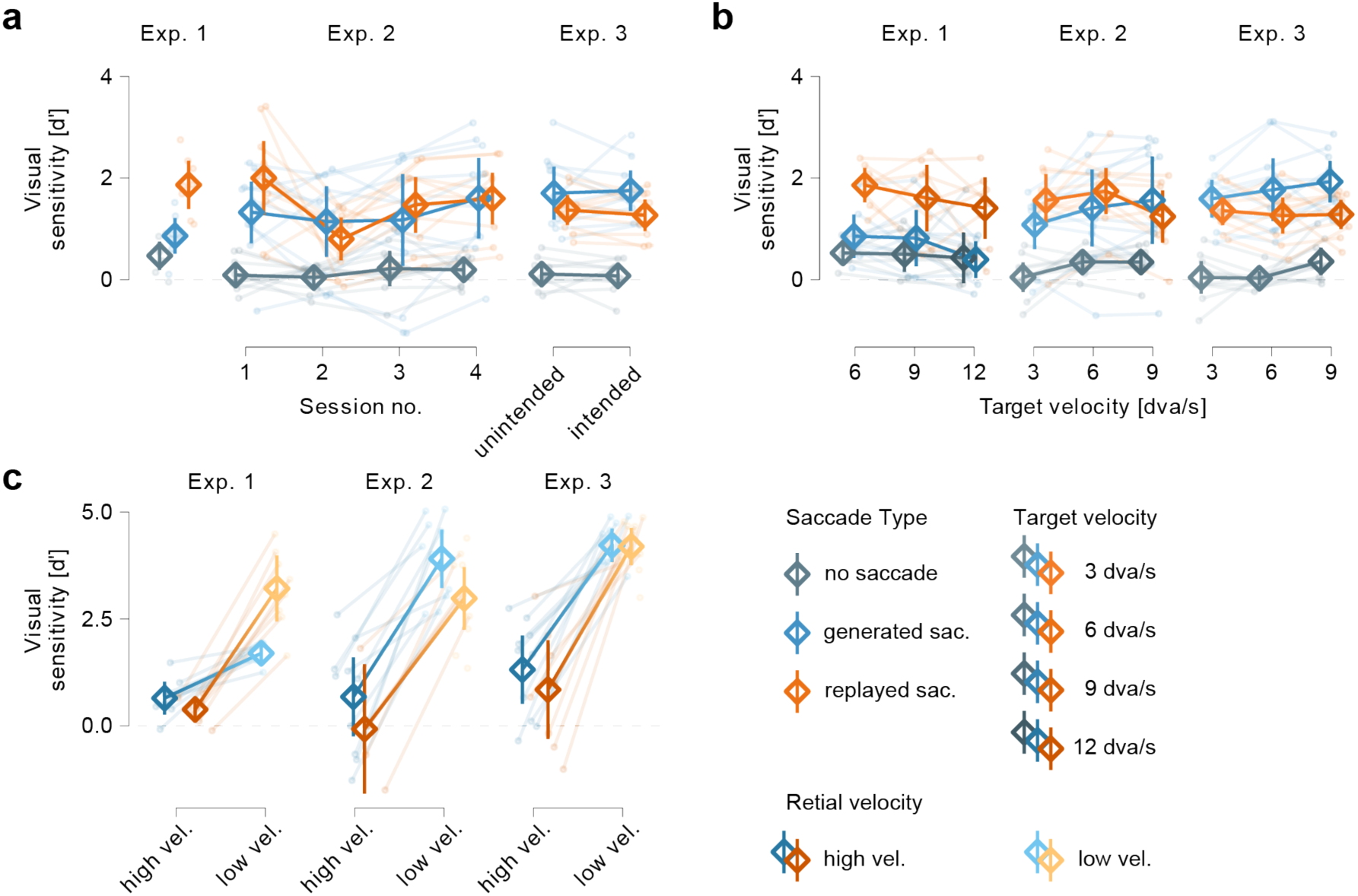
Visual sensitivity depends on saccade generation and their kinematics, but not on target velocity. **a** Visual sensitivity to the stimulus in trials without a catch-up saccade, and with either a generated or replayed catch-up saccade. Results are additionally plotted separately across sessions (**Exp. 2**) to assess whether the stimulus could have the intended training effect over time. Sensitivity is also shown by level of intention (**Exp. 3**), to determine whether explicitly instructing participants to either generate a saccade or maintain pursuit influenced stimulus perception. **b** Visual sensitivity plotted as a function of target velocity (3, 6, 9, and 12 dva/s, depending on the experiment). **c** Visual sensitivity as a function of retinal velocity, categorized into high (> 30 dva/s) and low (< 30 dva/s) velocity bins. Error bars represent 95% confidence intervals.

Across all experiments, visual sensitivity was substantially higher in trials with a generated or replayed catch-up saccade (**Exp. 1**: d’ = 1.38± 0.33, *p* < 0.001; **Exp. 2**: d’ = 1.55 ± 0.56, *p* < 0.001; **Exp. 3**: d’ = 1.43 ± 0.25, *p* < 0.001; **Fig. 2a**). To examine whether stimulus visibility was influenced by stimulus condition (generated vs. replayed saccade), session (**Exp. 2**), or instruction (**Exp. 3**), we conducted additional repeated-measures ANOVAs (rmANOVAs) for each experiment: In **Experiment 1**, a one-way rmANOVA revealed a significant main effect of stimulus condition (*F* (1,7) = 9.74, *p* = 0.017), with higher sensitivity for replayed (d’ = 1.87 ± 0.47) than for generated saccades (d’ = 0.86 ± 0.35; **Fig. 2a**)—likely reflecting a better match between phase shift velocity and replayed compared to generated saccade kinematics (see sections *Replay condition* in **METHOD DETAILS** and *Eye movement and stimulus parameters, visual sensitivity, and perception* in the **Discussion** for details on how (mis-)matching parameters likely affected visual sensitivity across experiments). In **Experiment 2**, a two-way rmANOVA found no significant difference in sensitivity between replayed (d’ = 1.47 ± 0.43) and generated (d’ = 1.31 ± 0.70) saccades (*F* (1,9) = 0.84, *p* > 0.250). However, there was a significant effect of session (*F* (3,27) = 4.56, *p* = 0.010) and a significant interaction between session and stimulus condition (*F* (3,27) = 6.58, *p* = 0.002; **Fig. 2a**), reflecting a notable drop in sensitivity in the second session, especially for replayed saccades. This decline likely reflects the effects of fine-tuning replay parameters to each observer’s eye movements, which improved visibility matching between generated and replayed saccades but reduced visibility in the replay condition. In **Experiment 3**, we found a significant effect of stimulus condition (*F* (1,9) = 6.46, *p* = 0.032), with slightly higher sensitivity for generated (d’ = 1.73 ± 0.41) than for replayed saccades (d’ = 1.32 ± 0.29; **Fig. 2a**). As expected, we found no significant effect of saccade type (intended vs. unintended; *F* (1,9) = 0.05, *p* > 0.250) and no interaction (*F* (1,9) = 1.05, *p* > 0.250).

We examined the effect of target velocity on stimulus visibility in separate analyses by calculating separate one-way rmANOVAs for every combination of experiment, saccade, and stimulus condition. Across experiments, target velocity did not significantly affect visual sensitivity in trials without saccadic eye movements (all *p*s > 0.066; **Fig. 2b**). However, we did find significant effects for generated saccades in **Experiment 1** (*F* (2,14) = 4.11, *p* = 0.039), and for replayed saccades in **Experiment 2** (*F* (2,16) = 4.30, *p* = 0.032; all remaining *p*s => 0.105; **Fig. 2b**). Because these isolated effects were neither consistent across experiments nor across stimulus conditions, we interpret them as unsystematic and most likely stemming from greater task demands at higher speeds (i.e., the additional effort required for the eyes to keep up with the fastest targets) rather than a genuine effect of velocity on stimulus visibility.

To determine whether visual sensitivity depended on the degree of retinal stabilization of the stimulus, we analyzed sensitivity based on retinal stimulus velocity. We focused on how closely the kinematics of generated or replayed saccades matched the stimulus parameters— assuming that a better match results in greater retinal stabilization. Retinal velocities were categorized as low (<30 dva/s) or high (>30 dva/s) depending on the combined velocity of eye movement and stimulus on the retina. We conducted individual two-way rmANOVAs for each experiment, with retinal velocity (low vs. high) and stimulus condition (generated vs. replayed) as factors, to assess whether the effect of retinal motion differed between eye movement types. In **Experiment 1**, we found a strong effect of retinal velocity (*F* (1,7) = 76.68, *p* < 0.001), with sensitivity being higher for low (d’ = 2.45 ± 0.45) than for high retinal velocities (d’ = 0.52 ± 0.25; **Fig. 2c**). We also observed a significant main effect of stimulus condition (*F* (1,7) = 8.04, *p* = 0.025), showing greater sensitivity for replayed (d’ = 1.80 ± 0.46) compared to generated saccades (d’ = 1.17 ± 0.23). Additionally, a significant interaction (*F* (1,7) = 107.1, *p* < 0.001) indicated that replayed saccades benefitted more from low retinal velocities than generated catch-up saccades (**Fig. 2c**). In **Experiment 2**, we again observed a strong effect of retinal velocity (*F* (1,4) = 56.16, *p* = 0.002), with no significant effect of stimulus condition (*F* (1,4) = 2.69, *p* = 0.177) and no interaction (*F* (1,4) = 0.04, *p* > 0.250), suggesting that visual sensitivity was primarily driven by the degree of retinal stabilization. **Experiment 3** showed a nearly identical pattern: a strong effect of retinal velocity (*F* (1,7) = 39.91, *p* < 0.001), in the absence of an effect of stimulus condition (*F* (1,7) = 0.91, *p* > 0.250), and no interaction (*F* (1,7) = 1.91, *p* = 0.210), confirming that low retinal velocity enhanced stimulus visibility.

Visual sensitivity in **Experiment 1** confirms that a mismatch between stimulus and saccade parameters decreases stimulus visibility and, hence, the effectiveness of saccade-contingent visual feedback. Results from **Experiments 2** and **3** further show that visual sensitivity depends not only on saccade generation but also on the degree of retinal stabilization provided by the saccade. Together, these findings demonstrate that the stimulus becomes visible saccade-contingently—whether generated or replayed—providing immediate visual feedback about saccade execution. It thus stands to reason that stimulus perception enabled participants to monitor their saccades and allowed us to isolate the influence of visibility on sensorimotor awareness during catch-up saccades in pursuit

### Motor control

To assess the extent of conscious control our observers exerted over catch-up saccade generation, we examined saccade rates (**Exp. 1**), their evolution over time while participants were instructed to suppress saccades (**Exp. 2**), and the influence of intention—manipulated via trial-by-trial instructions to either saccade or pursue (**Exp. 3**). Additionally, we assessed the effect of stimulus presence (present vs. absent) on eye movement control, reasoning that visible stimuli might provide saccade-contingent visual feedback supportive of suppressing (later) catch-up saccades. A two-way rmANOVA in **Experiment 1** revealed a significant main effect of target velocity (*F* (2,14) = 28.1, *p* < 0.001), with saccade rates increasing as target velocity increased (6 dva/s = 0.98 ± 0.33 s^-1^; 9 dva/s = 1.30 ± 0.42 s^-1^; 12 dva/s = 1.55 ± 0.53 s^-1^; **Fig. 3a**). Neither the main effect of stimulus presence (*F* (1,7) = 1.78, *p* = 0.224; **Fig. 3b**) nor the interaction between stimulus presence and velocity reached significance (*F* (2,14) = 3.17, *p* = 0.073), suggesting that the visibility of the stimulus did not facilitate suppression of catch-up saccades. A Bayesian model comparison, conducted to corroborate these null-results, provided very strong evidence for an effect of target velocity over the null model (BF10 = 1.0 × 10^9^), while the model with only stimulus presence was not supported (BF10 = 0.30). Including both main effects slightly reduced model evidence (BF10 = 5.0 × 10^8^) and adding their interaction further decreased support (BF = 0.31 relative to the model comprising only the two main effects), providing no evidence for an interaction between stimulus presence and target velocity.

**Fig 3.**
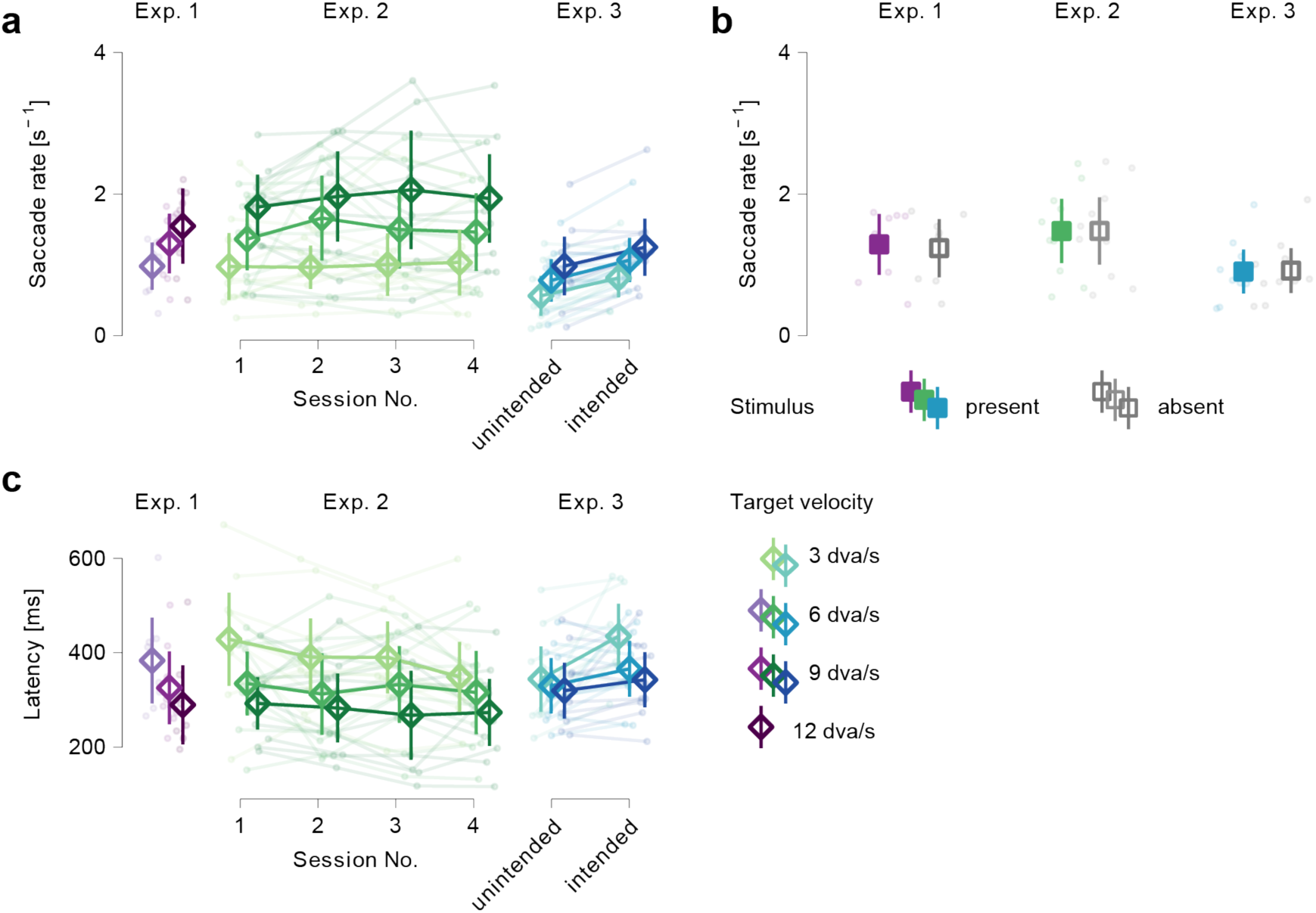
Saccade rate increases with target velocity in all experiments and decreases with explicit pursuit instruction (Exp. 3). **a** Saccade rate as a function of target velocity (all experiments), how it develops with training (e.g., session number; **Exp. 2**), and following the explicit instruction to pursue or saccade (e.g., unintended vs. intended, **Exp. 3**). **b** Saccade rate as a function of stimulus presentation for all experiments. **c** Saccade latency displayed using the same structure as in a. In all panels: Error bars represent 95% confidence intervals.

This finding was confirmed by the data from **Experiment 2**: We observed a comparable increase in saccade rate with target velocity (3 dva/s = 1.00 ± 0.18 s^-1^; 6 dva/s = 1.50 ± 0.23 s^-1^; 9 dva/s = 1.94 ± 0.28 s^-1^; **Fig. 3a**), which was confirmed as statistically significant by a three-way rmANOVA (*F* (2,16) = 22.42, *p* < 0.001). Neither the effect of stimulus presence (*F* (1,8) = 0.002, *p* > 0.250 **Fig. 3b**) nor that of session (*F* (3,24) = 0.64, *p* > 0.250) was significant, suggesting that stimulus presentation did not help participants suppress their catch-up saccades, nor did repeated exposure across sessions lead to a reduction in catch-up saccade generation over time. Additionally, none of the interactions were significant (all *p*s > 0.232). Bayesian model comparison yielded strong evidence for models including target velocity (BF10 = 1.2 × 10^29^), while models that excluded target velocity while including stimulus presence (BF10 < 0.14) or session (BF10 < 0.05) were not supported. Even the most complex interaction models were decisively less supported than the model solely including target velocity (BF = 0.14), providing no indication of added explanatory value by factors beyond target velocity alone. An analysis of potential long-term training effects comparing expert and naïve participants) is appendant to this manuscript (see **Supplementary Material** *S2: Observer groups in **Experiment 2***).

In **Experiment 3**, we again observed a significant effect of target velocity on saccade rate. Saccade rates increased significantly with increasing target speed (*F* (2,18) = 16.78, *p* = 0.003; 3 dva/s = 0.69 ± 0.19 s^-1^; 6 dva/s = 0.92 ± 0.21 s^-1^; 9 dva/s = 1.12 ± 0.27 s^-1^; **Fig. 3a**).

Importantly, our three-way rmANOVA revealed a significant difference between intended and unintended saccades (*F* (1,9) = 16.41, *p* = 0.003), indicating that participants generated significantly fewer saccades when explicitly instructed to pursue (mean = 0.77 ± 0.33 s^-1^) compared to when instructed to make a saccade (mean = 1.05 ± 0.32 s^-1^). Stimulus presence, on the other hand, did not significantly affect saccade rate (*F* (1,9) = 0.94, *p* > 0.250; **Fig. 3b**). This demonstrates again that saccade-contingent visual feedback did not facilitate conscious control over catch-up saccades. We also did not observe any significant interactions between other factors (all *p*s > 0.182). Like in the previous two experiments, Bayesian model comparison showed overwhelming evidence favoring models including target velocity (BF10 > 1.2 × 10^9^ for all such models). Unlike in the previous experiments, in **Experiment 3** we observed the highest Bayes factor for the model including saccade type as well as target velocity (BF10 = 7.57 × 10^18^). Models excluding target velocity or including only stimulus presence or saccade type had substantially lower support (BF10 < 1.2 × 10^9^). Adding interactions involving saccade type, stimulus presence, and target velocity consistently decreased model evidence, indicating no meaningful contribution of these interaction terms beyond the main effects of target velocity or saccade type. Details on how saccade distance, direction, and cue timing affected saccade rate and amplitude in the instructed saccade trials are provided in the Supplementary Material (see **Supplementary Material**, *S3: Instructed saccade conditions in **Experiment 3***).

We conducted a secondary analysis on the latency of the first saccade in each trial, hypothesizing that saccade frequency might be too coarse a measure to detect subtle effects of stimulus presence, training, or intention. We reasoned that participants might be able to delay—or “hold of” on—generating a saccade for longer if provided with visual feedback, through training, or by explicit instruction. Across three separate rmANOVAs, we consistently found significantly decreasing saccade latencies with increasing target speeds in all experiments (all *p*s < 0.002; **Fig. 3c**), indicating a faster need for catch-up saccades as target velocity rises. Stimulus presence was consistently non-significant (all *p*s > 0.226), as was training in **Experiment 2** (*p* > 0.250). However, saccade type in **Experiment 3** had a significant effect on saccade latency (*p* = 0.048), with a significant interaction between target velocity and instruction emerging only in this experiment (*p* = 0.004). While this may reflect that participants can more easily delay saccades at lower target speeds, we caution the reader to overinterpret our results without further data given the relatively high p-value. Bayesian model comparisons for each experiment helped further evaluate the contributions of target velocity (across all experiments) and session (**Exp. 2**) or saccade type (**Exp. 3**) on saccade latency. In **Experiment 1**, the model including only target velocity and participant received the strongest support (BF10 = 9.6 x 10^6^) with all models receiving substantially less support (BF < 0.28). In **Experiment 2**, the same model was again best supported (BF10 = 1.77 × 10^14^), with less evidence for the model that additionally included session (BF = 0.30). In **Experiment 3**, however, the comparison determined that the model including both target velocity and saccade type was best supported (BF10 > 7.7 × 10^7^), while the simpler models including only saccade type (BF10 = 1.9 × 10^3^) target velocity (BF10 = 9.93 × 10^2^) received substantially less support. These results suggest that while target velocity was a consistent predictor across experiments, additional variance was explained by session in **Experiment 2** and, even more substantially, by saccade type in **Experiment 3**.

Across all analyses, saccade rates consistently increased with target velocity but decreased when saccades were unintended, suggesting that our trial-by-trial instructions helped participants suppress saccades and that a certain level of control is possible. The absence of effects from stimulus feedback and training, however, demonstrates that low-level factors—such as speed and explicit instructions—drive this modulation rather than high-level conscious control. These findings were supported by our secondary analysis of saccade latencies, which showed a similar pattern across conditions.

### Sensorimotor awareness

We examined sensorimotor awareness of catch-up saccades during pursuit by analyzing observer’s saccade sensitivity: their ability to distinguish between trials with and without a catch-up saccade. To evaluate whether saccade-contingent visual feedback provided by our stimulus enhanced detection, we analyzed this separately for trials with and without the stimulus. In **Experiment 1**, saccade sensitivity was close to zero, whether the stimulus was present (d’ = −0.02 ± 0.50) or absent (d’ = −0.06 ± 0.27). A one-way rmANOVA revealed no significant effect of stimulus presence (*F* (1,7) = 0.08, *p* > 0.250; **Fig. 4a**). A Bayesian model comparison corroborated this result, with the model including stimulus presence receiving less support than the null model (BF10 = 0.43), indicating no evidence for an effect of stimulus presence. Similarly, in **Experiment 2**, we found sensitivity equally close to zero irrespective of stimulus presence (present: d’ = 0.00 ± 0.34; absent: d’ = −0.04 ± 0.37; **Fig. 4a**). A two-way rmANOVA, which included session number to assess potential training effects, revealed no significant main effects of stimulus presence (*F* (1,9) = 0.07, *p* > 0.250) or session (*F* (3,27) = 0.93, p > 0.250). Despite a significant interaction (*F* (3,27) = 4.43, *p* = 0.012), indicating that the effect of stimulus presence varied across sessions, we conclude from this analysis that neither saccade-contingent feedback nor training reliably improved detection performance. These findings were again corroborated by a Bayesian model comparison, which showed that all models—including those with stimulus presence or session—received less support than the null model (BF10 < 0.25). Sensitivity in **Experiment 3** was similarly low, with values near zero for both stimulus-present (d’ = 0.11 ± 0.24) and stimulus-absent trials (d’ = 0.16 ± 0.25; **Fig. 4a**). Finally, to determine whether movement intention (potentially combined with feedback) affected sensorimotor awareness of saccades, we conducted a two-way rmANOVA with stimulus presence and level of intention (intended vs. unintended) as factors. Neither the main effects (stimulus presence: (*F* (1,9) = 0.18, *p* > 0.250; saccade type *F* (1,9) = 0.00, p > 0.250) nor their interaction (*F* (1,9) = 1.87, p = 0.206; **Fig. 4a**) were significant, suggesting, once again, low saccade sensitivity that remained largely unaffected by feedback and movement intention. In Bayesian model comparison, all models received less support than the null model (BF10 < 0.34), regardless of whether they included stimulus presence, saccade type, or their interaction.

**Fig 4.**
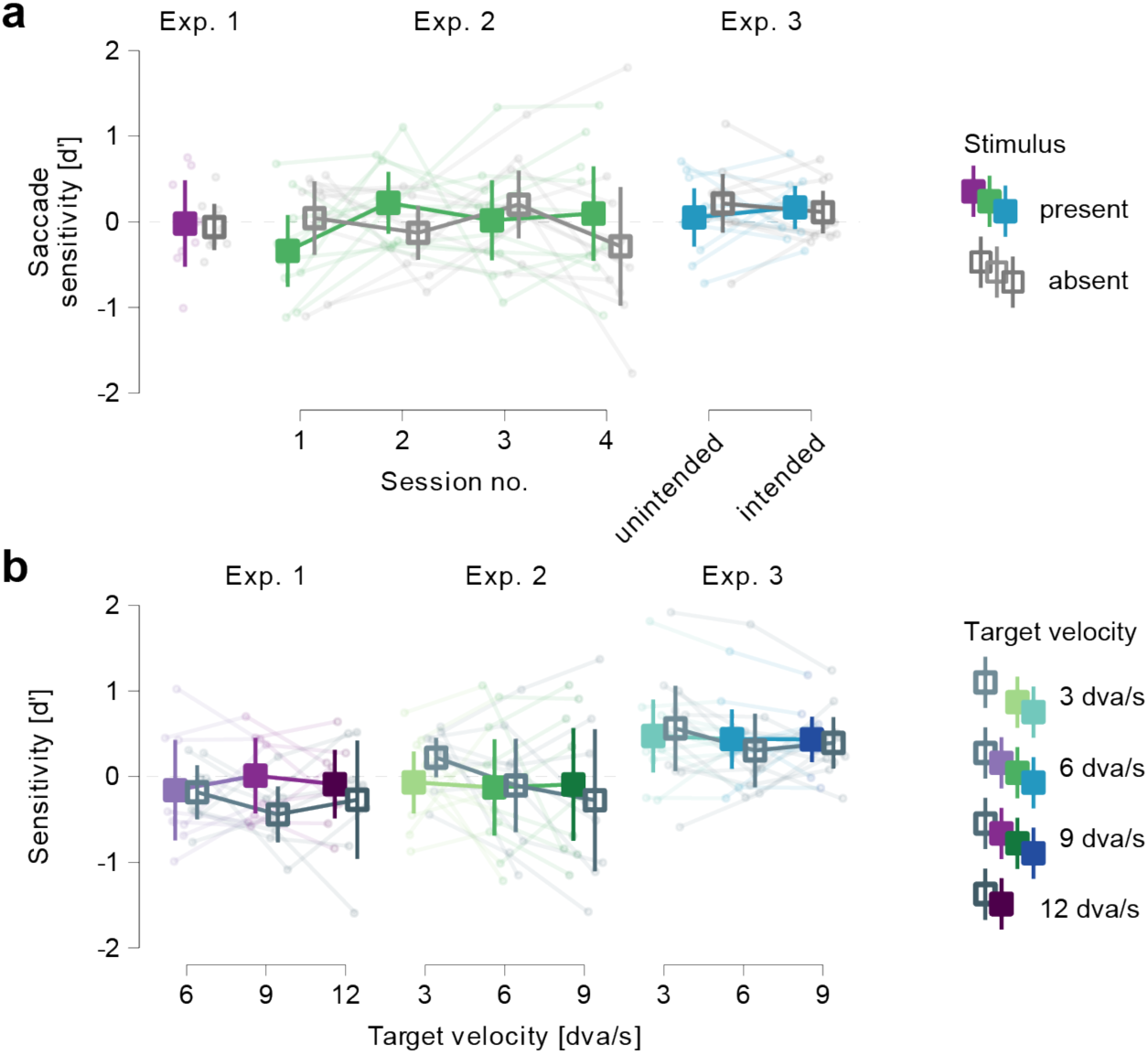
Low saccade sensitivity does not benefit from saccade-contingent feedback, cannot be trained (Exp. 2) and does not improve when movements are explicitly instructed (Exp. 3). **a** Saccade sensitivity as a function of stimulus presence (all experiments), its development over time (e.g., session number; **Exp. 2**), and following the instruction to pursue or make a catch-up saccade (e.g., unintended vs. intended, **Exp. 3**). **b** Saccade sensitivity as a function of stimulus presentation for all experiments and target velocities. In all panels: Error bars represent 95% confidence intervals.

To examine whether target velocity influenced saccade sensitivity, we conducted separate two-way rmANOVAs for each experiment, with stimulus presence (absent vs. present) and target velocity as factors (6, 9, 12 dva/s in **Exp. 1**; 3, 6, 9 dva/s in **Exp. 2** and **3**). Across all experiments, we found no significant main effects or interactions (all *p*s > 0.160, BF10 < 0.85), indicating that variations in target velocity did not affect saccade detection performance, regardless of stimulus presence (**Fig. 4b**). To compare saccade sensitivity across experiments, we conducted a one-way ANOVA with experiment as the sole between-subjects factor. Although the analysis revealed no significant differences between experiments (*F* (2,25) = 1.92, *p* = 0.168, BF10 = 0.70), descriptive statistics showed that saccade sensitivity—while generally low—was significantly above zero in **Experiment 3** (**Exp. 3**: d’ = 0.45 ± 0.29), compared to non-significant values in the other two experiments (**Exp. 1**: d’ = 0.02 ± 0.39; **Exp. 2**: d’ = 0.45 ± 0.47; **Fig. 4b**). Note that we averaged over trials with and without saccade-contingent visual information in this final analysis. The higher saccade sensitivity observed here compared to other analyses suggests that participants may have been leveraging the visual cues provided by actually seeing the stimulus. Nevertheless, the comparatively higher saccade sensitivity in **Experiment 3** suggests that intentional engagement may have played a role in modulating saccade awareness, even though overall differences between experiments were not statistically significant.

Across all analyses, saccade sensitivity remained consistently low, unaffected by stimulus presence, target velocity, training (**Exp. 2**), or movement intention (**Exp. 3**). Visual feedback alone did not enhance detection performance. Notably, while movement intention has been shown to influence microsaccade awareness (Klanke et al., 2025), and successfully modulated voluntary control over catch-up saccades in the current study, it did not improve sensorimotor awareness. These findings suggest that awareness of catch-up saccades during pursuit is minimal and resistant to both perceptual and intentional modulation.

## Discussion

### Voluntary control of catch-up saccades

In a series of three experiments, we examined whether observers could control catch-up saccades during pursuit and whether they were aware of these brief ballistic eye movements. While participants were able to exert some voluntary control over their catch-up saccades, as indicated by reduced rates when explicitly (i.e., visually) prompted to suppress them, this ability was markedly limited: control did not improve with training (**Fig. 3a**), and participants were unable to use immediate saccade-contingent visual feedback to further reduce saccade rates (**Fig. 3b**). Instead, saccade rate was most consistently modulated by target velocity, with more saccades occurring at higher speeds (**Fig. 3a**). This pattern indicates that catch-up saccade generation is primarily driven by task demands—potentially to maintain foveation on a fast-moving target (c.f., Heinen et al., 2016) or to correct for low-level position errors (i.e., ‘retinal slip’; Daye et al., 2014; Mcilreavy et al., 2019; Schröder et al., 2023)—rather than conscious control. Interestingly, the reduction in saccade rates following the suppression cue may similarly reflect the influence of task dynamics rather than the prompt itself. By presenting this cue visually on every trial, we likely allowed participants to adjust their behavior indirectly, responding to the visual information inherent in the task rather than through a direct exertion of will. Further support for this interpretation comes from our analysis of intended catch-up saccades in **Experiment 3**. We found that participants adjusted their saccade rates primarily based on temporal aspects of the task, making more saccades when the go-cue appeared earlier, and only to a lesser extent in response to spatial factors. In contrast, they modulated saccade amplitude mainly according to spatial features—such as the distance and direction of the instructed saccade—while cue timing had little to no effect (see **Supplementary Material** *S3: Saccade parameters in response to target manipulations in **Experiment 3***). These results suggest that participants adapted their eye movements in response to the visually presented goal (i.e., the target position), corroborating the idea that saccades can be modulated voluntarily during pursuit. Together, these findings demonstrate that low-level sensorimotor factors primarily drive catch-up saccade generation during pursuit and suggest that while conscious, top-down control over these movements is possible, it remains limited.

### Sensorimotor awareness of catch-up saccades

While participants in our study were able to exert some volitional control over their catch-up saccades, this control did not lead to increased sensorimotor awareness. Across all experiments, awareness of these saccades remained low, regardless of stimulus presence, training, movement intention (**Fig. 4a**), or eye-movement expertise (see **Supplementary Material** *S2: Observer groups in **Experiment 2***). This dissociation suggests that although catch-up saccades can be modulated intentionally to some extent, they remain largely inaccessible to conscious monitoring, pointing to a functional separation between oculomotor control and introspective access. This is particularly surprising given the comparatively large amplitudes of the catch-up saccades in our data—which, depending on the experiment and condition, averaged around 1.5 dva. In contrast, a study by Klanke et al. (2025), which investigated sensorimotor awareness of microsaccades, found that these much smaller eye movements—only 1 dva or less—were nonetheless sometimes accessible to introspection. A possible explanation for the low sensorimotor awareness of catch-up saccades in the present study is that participants’ attention was focused on the ongoing smooth pursuit—potentially masking awareness of saccades embedded within it. In this view, the seamless nature of pursuit may create an illusion of uninterrupted tracking, rendering discrete corrective movements like catch-up saccades introspectively invisible. This aligns with the idea that the initiation and control of pursuit eye movements require considerable attentional resources (Chen et al., 2002; Kerzel et al., 2009; Khurana & Kowler, 1987).

The effortlessness with which catch-up saccades are usually blended into our sense of a fluid, continuous pursuit becomes most apparent when the (predictive) pursuit machinery breaks down. Koerfer, Watson & Lappe (2024) provide a vivid demonstration with their non-rigid moving vortex: during fixation the pattern looks perfectly coherent, yet when observers try to track it, smooth pursuit gain collapses to almost zero and the target can be followed only by a string of catch-up saccades (Koerfer et al., 2024). The attempt to pursue the stimulus thus turns the normally imperceptible catch-up saccades into conspicuous sensorimotor events, with the disruption of perceptual flow allowing awareness of each corrective eye movement.

Interestingly, our data contrasts somewhat with recent findings by Goettker et al. (2024), who reported that observers were able to evaluate the accuracy of their combined pursuit and saccadic eye movements when tracking unpredictable targets. However, several differences between the paradigms may help reconcile these results. For one, the task in Goettker et al. emphasized tracking accuracy rather than awareness of movement occurrence, potentially engaging different cognitive processes. Additionally, their paradigm utilized visual information and performance history (e.g., gaze–target deviation from a visible sinusoidal trajectory and self-comparison to past performance), potentially allowing participants to rely on external visual cues and performance heuristics, rather than direct introspective access to eye movements themselves. Finally, even in Goettker et al.’s study, metacognitive sensitivity for eye movements remained considerably lower than for hand movements, reinforcing the idea that access to oculomotor events is fundamentally constrained—even when conditions favor introspective awareness. Overall, our findings complement those of Goettker et al. by highlighting that, even when some degree of access to catch-up saccades is possible—as their results suggest—conscious awareness of these movements remains limited, particularly when external cues and comparative feedback are minimized.

### Eye movement and stimulus parameters, visual sensitivity, and perception

Our analysis of visual sensitivity revealed that stimuli with rapid temporal phase shifts can selectively target specific eye movement types—such as catch-up saccades—while remaining largely invisible during others, like smooth pursuit (**Fig. 2a**/**b**). Crucial to this selective visibility is a precise alignment between the stimulus properties and the dynamic parameters of the eye movements. Even slight mismatches in frequency, velocity, or timing—whether in the stimulus design or assumptions about the eye movements—can substantially reduce stimulus visibility and thus diminish its effectiveness. In **Experiment 1**, we assumed a relatively large and fast catch-up saccade profile when designing the stimulus, which led to a mismatch for many participants and allowed the stimulus to become faintly visible even during pursuit-only trials. In contrast, fine-tuning the stimulus to better match individual saccade dynamics in **Experiments 2** and **3** effectively eliminated visibility in the absence of saccades and optimized contingent visibility during catch-up saccades. Across all experiments, visual sensitivity was closely tied to retinal stabilization, with low retinal velocities consistently producing greater sensitivity. Our data hence demonstrate the importance of calibrating saccade-contingent stimuli to individual eye movement characteristics and emphasize the role of fine-grained sensorimotor tuning in shaping visual perception during movement.

Seeing the stimulus saccade-contingently does not necessarily imply that participants understood the systematic relationship between their eye movements and the perceptual feedback. In **Experiment 1**, we additionally asked participants how certain they were that a catch-up saccade had caused the stimulus to become visible—if they had previously reported both seeing the stimulus and making a saccade—or, alternatively, how certain they were that the stimulus had not been caused by a saccade—if they reported seeing the stimulus but denied making an eye movement. Our analysis of these responses suggests generally low certainty, with answers clustering around the midpoint of the scale (i.e., the point of highest uncertainty; see **Fig. S1**). This indicates that participants were, on average, unable to reliably distinguish between trials in which the stimulus was caused by a saccade and those in which it was not. While this could in part be driven by the suboptimal stimulus configuration in **Experiment 1**, we believe it primarily reflects a general ambiguity regarding the connection between saccades and their visual consequences—particularly when alternative perceptual interpretations (i.e., an identical visual event occurring without a saccade, as in the replay condition) are presented alongside the saccade-contingent change in visual perception (see section *S1: Causal assignment from **Experiment 1*** in the **Supplementary Material**).

### The role of intention for sensorimotor awareness

In **Experiment 3**, our goal was to manipulate movement intention by instructing participants either to pursue the target naturally (unintended saccade condition) or to generate a catch-up saccade deliberately (intended saccade condition). It remains an open question whether this truly reflects a change in intention as opposed to a strategic response to task demands or an effect of attention. However, the robust increase in saccade rates in the intended saccade condition compared to the unintended one suggests that the manipulation successfully altered participants’ volitional engagement with their eye movements. Surprisingly, despite this intentional engagement, saccade sensitivity—that is, participants’ awareness of their own catch-up saccades—remained very low, especially in light of recent findings by Klanke et al., (2025) who reported higher awareness for microsaccades under similar conditions. A supplementary analysis revealed that this disconnect was due to a significant increase in both hit and false alarm rates when saccades were instructed, suggesting that while participants were more responsive overall, they were not more accurate in distinguishing when a saccade had actually occurred (see **Supplementary Material** *S4: A closer look at saccade sensitivity: Hit and false alarm rates across experiments*). Crucially, our data therefore support that our manipulation indeed affected intention rather than simply task strategy or attention: Participants not only followed the instruction to make a saccade as well as they could, but also genuinely believed they had done so—even when they had not. To better understand how intention influenced awareness, we compared saccade sensitivity across all three experiments. Our analysis revealed that while sensitivity was slightly above zero in all three experiments, it was only significantly different from zero in **Experiment 3**. This aligns with Klanke et al.’s (2025) finding that intention can enhance awareness for microsaccades, irrespective of whether the microsaccades were intended or unintended. Notably, **Experiment 3** included both intended and unintended saccade conditions, whereas the other experiments—particularly **Experiment 1**, which showed the lowest saccade sensitivity— treated catch-up saccades as spontaneous. The slight increase in awareness observed in **Experiment 3** suggests that explicit intention can moderately enhance sensorimotor sensitivity, even though overall awareness remains low. While our manipulation of intention was, hence, likely successful, its effect on saccade awareness was minimal, indicating that conscious access to saccades during ongoing pursuit remains limited even when these movements are voluntarily produced.

### In pursuit of saccade awareness

Smooth pursuit has long been understood as a voluntary eye movement (c.f., Kowler, 2011) that involves both sensory inputs and cognitive influences. Especially Eileen Kowler’s influential research highlighted how pursuit can be modulated by cognitive processes like attention (Khurana & Kowler, 1987; Murphy et al., 1975), expectation (Kowler, 1989; Kowler et al., 1984; Kowler & Steinman, 1979, 1981), and learning (Kowler, 1989, 2011; Kowler et al., 1984)—in addition to task affordances (Kowler & McKee, 1987). Our findings complement the work by Eileen Kowler and extend it to catch-up saccades. Although our data show that these corrective saccades are predominantly shaped by low-level task demands such as target velocity, we also observed a small but reliable effect of intention on saccade generation— suggesting that motor control of saccades during pursuit is open to top-down modulation and responsive to cognitive influences. While Eileen Kowler did not explicitly address conscious awareness of pursuit eye movements in her research, her seminal findings on anticipatory pursuit support the idea that observers consciously generate and are aware of these movements. In stark contrast, our research indicates that catch-up saccades almost always escape awareness. While intention may modestly enhance awareness (see previous section: *The role of intention for sensorimotor awareness*), we consistently found saccade sensitivity near zero across all experiments and conditions. This suggests that, perhaps unlike smooth pursuit, catch-up saccades remain largely inaccessible to conscious monitoring. Together, our findings suggest that while pursuit eye movements and catch-up saccades are tightly linked components of oculomotor behavior, they differ fundamentally in how they interface with voluntary control and awareness. Our work extends Eileen Kowler’s research by revealing that voluntary control does not necessarily extend to awareness, even for closely linked eye movement behaviors.

## Conclusion

When we watch koi in a pond, we experience the illusion that they remain at the center of our gaze despite their constant slow coasting. This illusion persists even though our smooth pursuit is far from perfect and is frequently interrupted by catch-up saccades—even when the gaze target moves at low speeds. Our data suggest that catch-up saccade frequency is strongly modulated by target velocity, with more saccades occurring at higher speeds. These saccades are open to conscious motor control—if the observer can exploit dynamic visual information to modulate their eye movements—but remain inaccessible to introspective awareness, even when accompanied by (trans-saccadic) visual transients. This dissociation between control and awareness highlights the reflexive, opaque nature of corrective eye movements and suggests that the visual system favors visual stability over introspective access to the eye movements that enable it: We can follow the koi effortlessly—without ever noticing the corrections our eyes perform along the way.

## Acknowledgements

JNK was supported by the Berlin School of Mind and Brain, Humboldt-Universität zu Berlin. SO was supported by the DFG (OH 274/4-1) and funding from the Heisenberg Programme of the DFG (OH 274/5-1). MR was supported by the European Research Council (ERC) under the EU’s Horizon 2020 research and innovation program (grant agreement no. 865715), and by the DFG (grants RO3579/8-1 and RO3579/10-1). We also thank Alexis Soucy for collecting the data for Experiment 2.

## STAR Methods

### RESOURCE AVAILABILITY

#### Lead contact

Information and requests regarding resources for this study should be directed to and will be fulfilled by the lead contact, Jan-Nikolas Klanke

### Materials availability

There are no restrictions for the distribution of materials.

### Data and code availability

- The preregistration, data, and analysis code for **Experiment 1** has been deposited at the Open Science Framework and will be made publicly available as of the date of publication. [LINK WILL FOLLOW HERE UPON PUBLICATION].
- The preregistration, data, and analysis code for **Experiment 2** has been deposited at the Open Science Framework and will be made publicly available as of the date of publication. [LINK WILL FOLLOW HERE UPON PUBLICATION].
- The preregistration, data, and analysis code for **Experiment 3** has been deposited at the Open Science Framework and will be made publicly available as of the date of publication. [LINK WILL FOLLOW HERE UPON PUBLICATION].

### EXPERIMENTAL MODEL AND SUBJECT DETAILS

In **Experiment 1**, a total of 8 participants were recruited by means of the “Psychologischer Experimental-Server Adlershof” (PESA) of the Humboldt-Universität zu Berlin. Participants (4 female, 0 diverse) had a mean age of 25 years (*SD =* 3.9, *min* = 21, *max* = 33). Of our participants, 7 were right-handed and one was left-handed. Similarly, 7 were right-eye dominant, and one participant was left-eye dominant. All 8 participants had normal or corrected-to-normal vision.

In **Experiment 2**, a total of 10 participants were recruited by means of the “Psychologischer Experimental-Server Adlershof” (PESA) of the Humboldt-Universität zu Berlin and from members of the laboratory. Participants (9 female, 0 diverse) had a mean age of 25.5 years old (*SD =* 3.3, *min* = 21, *max* = 30). Of these participants, 9 were right-handed and one was left-handed. Similarly, 9 participants were right-eye dominant, one participant was left-eye dominant. All 10 participants had normal or corrected-to-normal vision.

In **Experiment 3**, a total of 10 participants were recruited by means of the “Psychologischer Experimental-Server Adlershof” (PESA) of the Humboldt-Universität zu Berlin. Participants (7 female, 1 diverse) had a mean age of 22.7 years old (*SD* = 2.1, *min* = 20, *max* = 25), and all 10 were right-handed and 6 were right-eye dominant. All ten participants had normal or corrected-to-normal vision. Participants were paid upon completion of the last session. The compensation was based on an hourly rate of €10/hour. Alternatively, psychology students could choose to obtain participation credit (1 credit per 15 minutes of participation) required for the successful completion of their bachelors’ program.

Participants in all three experiments were paid upon completion of the last session. The compensation was based on an hourly rate of €10/hour. Alternatively, psychology students could choose to obtain participation credit (1 credit per 15 minutes of participation) required for the successful completion of their bachelors’ program.

#### Exclusion of participants

For **Experiment 1, 2** and **3**, we pre-registered an exclusion criterion that ensured that participants would not participate if they showed the inability to execute stable fixation or correct eye movements: The inability to complete at least 4 blocks during the first experimental session due to fixation failures led to immediate exclusion from the experiment in all experiments.

In **Experiment 1**, no participants were excluded from data collection; however, one participant chose to discontinue their participation after completing the first session for personal reasons. In **Experiment 2**, one participant was excluded due to an eye tracker malfunction that occurred during the fifth block of the first session, resulting in no data being saved for the entire session. To minimize the impact of missing data, we discontinued the experiment for this participant. Additionally, two other participants withdrew after partially completing the first session for personal reasons. In **Experiment 3**, no participants were excluded, but one chose to discontinue after completing two of the four sessions for personal reasons.

In all experiments, data collection continued until the full pre-registered sample size was reached: 8 participants for **Exp. 1**, and 10 for **Exp. 2** and **3**.

### METHOD DETAILS

#### Apparatus

Participants were seated in a dark room in front of a screen at a distance of 340 cm and their head stabilized using a chin rest. We projected visual stimuli on a 141.0 x 250.2 cm video-projection screen (Stewart Silver 5D Deluxe; Stewart Filmscreen, Torrance, CA, USA) using a PROPixx DLP (960 × 540 pixels; VPixx Technologies Inc., Saint Bruno, QC, Canada) with a refresh rate of 1440 Hz. We recorded participants’ eye positions of both eyes with a head-mounted eye tracker at a sampling rate of 500 Hz (EyeLink 2 Head Mount; SR Research, Ottawa, ON, Canada). The experiments were controlled on a workstation running the Debian 8 operating system, using Matlab (Mathworks, Natick, MA), the Psychophysics Toolbox 3 (Brainard, 1997; Kleiner et al., 2007; Pelli, 1997) and the EyeLink Toolbox (Cornelissen et al., 2002).

#### Eye movement task and Rashbass’ paradigm

To examine control and awareness of catch-up saccades during pursuit, we employed a version of the Rashbass paradigm (Rashbass, 1961), which minimizes initial catch-up saccades by allowing the visual system to prepare a pursuit response before target onset. In our adaptation, used in **Experiments 1** and **2**, participants tracked a black target (0.35 dva diameter) moving in a straight horizontal line across the screen midline; target velocities were 6, 9, or 12 dva/s in **Experiment 1** and 3, 6, or 9 dva/s in **Experiment 2**, corresponding to movement amplitudes of 6, 9, or 12 dva (**Exp. 1**) and 3, 6, or 9 dva (**Exp. 2**), respectively. To facilitate pursuit initiation without early saccades, each trial began with a fixation interval during which participants maintained gaze on a central fixation dot while the moving target—initially offset by 1.5, 3, or 4.5 dva—was already in motion toward the fixation point. This setup ensured that the pursuit target crossed the fixation location at the moment pursuit was to begin, enabling smooth tracking without the need for a corrective catch-up saccade. Participants were instructed to maintain fixation until the target reached the fixation location, after which they were to pursue the target smoothly. Target motion was continuous throughout the trial, with no pauses or halts. Target velocity was blocked, and participants were informed before each block whether target speed would be low, medium, or high.

In **Experiment 3**, we extended this approach by using more visually complex pursuit targets to test eye movement awareness under increased perceptual demands. Instead of a single moving dot, the pursuit target consisted of a cloud of 4 to 8 black dots (each 0.22 dva in diameter), with dot positions sampled from a 2D Gaussian distribution (x: *M* = 0, *SD* = 1 dva; y: *M* = 0, *SD* = 0.4 dva), resulting in a horizontally elongated shape. Targets moved horizontally across the screen midline—either left to right or right to left—at constant speeds of 3, 6, or 9 dva/s (same as in **Exp. 2**), covering distances of 3, 6, or 9 dva, respectively. As in **Experiments 1** and **2**, a fixation interval preceded the pursuit period: participants maintained fixation on a central dot while the moving target was already visible and approached the fixation point from an initial offset (1.5, 3, or 4.5 dva), resulting in total target amplitudes of 4.5, 9, and 13.5 dva. Participants were instructed to pursue the target only once it reached the fixation location. Target motion was continuous, and target velocity was blocked in random order and announced prior to each block.

To investigate the role of intention in saccade awareness during pursuit, each trial in **Experiment 3** featured one of two eye movement instructions. In unintended saccade trials, participants were instructed to pursue the target as smoothly as possible; here, both the fixation point and all target dots were black, and any catch-up saccades were reflexive. In contrast, intended saccade trials required participants to deliberately generate a saccade during pursuit. These trials began with a white fixation dot that disappeared as soon as it was occluded by one of the moving target dots, which in turn changed to white to indicate the dot to be pursued. This dot remained white for 200–450 ms (early jump) or 700–950 ms (late jump) relative to pursuit onset, after which a second target dot turned white for 50 ms to signal the saccade target. All dots then returned to black, indicating that the participant should execute the instructed saccade. Jump targets were offset by 0.5, 1, or 1.5 dva horizontally (left or right of the pursuit target) and included a vertical offset of ±5° to introduce oblique saccades and reduce trial predictability. The early and late jump conditions were designed to manipulate participants’ ability to comply with the saccade instruction, with early jumps facilitating and late jumps potentially hindering timely execution.

#### The stimulus

The stimulus was formed by two vertically oriented sinusoidal gratings with spatial frequency of 5 cpd, combined with identical cosine-tapered masks. The combined gratings and masks appeared like two striped horizontal bars that smoothly blended in with the gray background (see **Fig. 1b and d**). Both stimuli had a length of 28 dva and height of 2 dva. The size of the tapered sections is always 1. Masks were created by generating separate cosine tapered windows for the height and the width of each stimulus that are then combined by multiplication (again, separately per each stimulus).

To examine if a visual consequence affected awareness of the underlying eye movement (**Exp. 1**, **2**, and **3**), our stimulus was designed to be invisible during pursuit, but visible when briefly stabilized on the retina by a saccade. To achieve invisibility, we added a high-velocity phase shift to the grating, creating a temporal frequency above 60 Hz (cf. Castet & Masson, 2000). In **Experiment 1**, phase shift was based on the peak velocity of a saccade with an amplitude of 2 dva (i.e., 100.65 dva/s according to the formula and values reported by Collewijn et al., 1988). We added the speed of the pursuit target the phase shift velocity to ensure invisibility in the absence of a saccade, leading to phase shift velocities of 106.65, 109.65, and 112.65 dva/s respectively. Because of a poor stimulus visibility in the first experiment and the lower pursuit target speeds, we decided to use a lower phase shift velocity in **Experiment 2** and **3.** There, the phase shift velocity was based on a saccade with an amplitude of 1 dva, resulting in a base velocity of 53.50 dva/s. After correcting for the speed of the pursuit target, the phase shift velocity was set to be 56.50, 59.50, or 62.50 dva/. Presented in this way, the stimulus should become visible only during catch-up saccades, which transiently reduce the retinal velocity of the grating. This momentary stabilization can raise the grating above the detection threshold, much like how saccades have been shown to reveal otherwise invisible high-frequency patterns (Deubel et al., 1987; Deubel & Elsner, 1986; Kelly, 1990).

The direction of the phase shift—solely determined by the stimulus orientation—can be either rightward (stimulus orientation of 0 deg from vertical) or leftward (stimulus orientation of 180 deg from vertical) to match the direction of potential catch-up saccades (the smooth pursuit target will be moving purely horizontally). In **Experiment 1**, phase shifts were always oriented in the same direction. To increase visibility of the stimulus, the stimulus’ phase shifts were oriented in opposite directions in **Experiment 2** and **3**. Stimulus presentation started at a contrast of 0 and was ramped up to maximum contrast of 50% within 200 ms. Conversely, in the last 200 ms of presentation, the stimulus contrast slowly decreased, and the stimulus faded out. The stimulus was thusly modulated to avoid sudden onsets and offsets that might have led to transient changes in stimulus visibility. Because generated catch-up saccades rendered it visible, this stimulus condition was called *generated saccade* condition throughout **Experiment 1**, **2** and **3**.

#### Replay condition

We compared stimulus visibility for generated saccades with two other conditions: In one condition, we used the same stimulus (as described in the previous paragraph) in terms of its spatial frequency, size, the extent of its tapered section, its orientation, contrast modulation, phase shift, as well as phase onset, placement, and presentation duration. We added a rapid change in the onscreen locations of the stimulus apertures to replicate the retinal consequence of a typical eye movement for the observer. If the aperture movement was in the opposite direction of the stimulus’ phase shift, the image of the stimulus appeared to slow down on the retina, resulting in retinal effects very similar to those of an actual catch-up saccade. The aperture motion was generated based on a fixed set of parameters and simulated a catch-up saccade of 2 dva and a peak velocity of 100.65 dva/s in all three sessions of **Experiment 1**. In the first session of **Experiment 2** and **3**, we used a similar approach for the first session and used the parameters of a saccade with an amplitude of 1 dva and a peak velocity of 53.5 dva (to match the lower velocity of the phase shift of the stimuli). In the later sessions, we adapted these parameters to the eye movement data of each participant to better match the stimulus visibility for generated catch-up saccades: we fit gamma functions to the distribution of saccade amplitudes measured for the three different target speed conditions (i.e., 3, 6, and 9 dva/s). We additionally fitted main sequence functions to the saccade amplitude and peak velocity data to determine the optimal peak velocity at any given amplitude for each observer (the parameters and formula for calculating the velocity profile of these simulated catch-up saccades were based on Collewijn et al., 1988). During the experiment, we sampled individual saccade amplitudes from observer-specific gamma distributions and determined corresponding peak velocities based on the main sequence fits. These parameters were then used to simulate biologically plausible, observer-specific catch-up saccade profiles, which were replayed via aperture movements. The direction of these simulated saccades was constrained to fall within ±2° of the horizontal axis (i.e., 0° or 180°).

Finally, in *no-stimulus condition* trials, the stimulus will be presented at 0% contrast while everything else will be identical to trials with generated and replayed saccades.

In **Experiment 1**, 20% of all trials were no-stimulus condition trials while the remaining 80% of trials will be with stimulus. Stimulus trials were split evenly between generated (40%) and replayed saccades (40%). To increase stimulus visibility and make it a more reliable predictor of saccade generation, we increased of number of generated saccade trials to 60% in **Experiment 2**. The remaining trials were split between the replayed saccade (20%) and no-stimulus condition (20%). Finally, in **Experiment 3**, we returned to a more even split between conditions and presented the stimulus 37.5% of trials in the generated saccade condition, 37.5% in the replayed saccade condition, and in 25% of trials in the no-stimulus condition.

### General Methods Experiment 1

#### Fixation-check interval

Before the start of each trial, a target-shaped central fixation dot appeared before an otherwise gray background. The fixation point (inner part) had a diameter of 0.2 dva while the outer ring had a diameter of 0.6 dva. Before the onset of each trial, a fixation control routine was run that required the gaze position of the observer to be inside a circular region (3 dva in diameter) centered on the fixation spot. The trial started only when the fixation control was successful for at least 100 ms.

#### Fixation interval

The start of the fixation interval was marked by the disappearance of the outer ring of the fixation point. Participants were instructed to look at the fixation dot and not move their eyes for the entire duration of this phase of the trial (500 ms). At the same time, a pursuit target appeared at either 3, 4.5, or 6 dva relative to the fixation dot, positioned towards the outer screen edge (opposite to the motion direction). It moved with constant speed of 6, 9, or 12 dva/s towards the fixation dot. The target was a black dot with a diameter of 0.35 dva. Participants were instructed to keep fixating until the target reached the fixation dot.

#### Pursuit and stimulus presentation interval

The pursuit and stimulus presentation interval began once the pursuit target fully occluded the fixation dot (i.e., the observer’s gaze position). Participants were instructed to start pursuing the target with their eyes when this occurred. In active or replay condition trials, two stimuli were presented 3 dva above and below the screen’s horizontal midline. The bands had a length of 28 dva and a height of 2 dva. The pursuit and stimulus presentation interval lasted for 1000 ms in total. Between this and the response interval, there was a short delay of 50 ms during which nothing was presented onscreen.

### Response interval

In the response interval, participants were always presented with two simple yes-no questions Firstly, we asked participants if they perceived the stimulus in the previous trial. We presented the question “Did you perceive a STIMULUS FLASH?”, together with the two response options “Yes!” and “No!”. In a second step, participants reported if they generated a catch-up saccade. We displayed the question “Do you think you generated a CATCH-UP SACCADE?” and the same response options “Yes!” and “No!”. Both questions could be answered by pressing the arrow key corresponding to the direction of the chosen response option (e.g., the right arrow key for a selected of the right-ward response prompt).

Participants’ responses to these first two questions determined the presentation of the final stage of the response phase—the link between eye movement and stimulus visibility: If participants reported that they perceived a stimulus flash and that they think they generated an eye movement, they were asked: “How sure are you that the stimulus WAS caused by a catch-up saccade?”. If they reported that they did not perceive the stimulus, but thought they generated an eye movement, we asked “How sure are you that the stimulus flash was NOT caused by a catch-up saccade?”. To answer, participants had to choose one of four options displayed on a continuous scale: “not sure”, “rather not sure”, “rather sure”, and “very sure”. Participants chose their response by adjusting the position of a response prompt via the arrow keys and submitted their response by pressing the space bar.

#### Variations in Experiments 2

We used the same procedure in **Experiment 2** as in the first experiment with only minor variations: Target speeds were lowered to 3, 6, and 9 dva/s due to the high number of catch-up saccades in all target speed conditions of **Experiment 1**. Initial target positions were, therefore, adjusted to 1.5, 3, and 4.5 dva relative to the fixation dot. Because, unlike in the first experiment, the phase shift of the two stimuli bands was always in opposite directions in **Experiment 2**, we added a simple localization task to the response interval: If participants reported that they perceived the stimulus in response to the first question, we asked if they perceived the one above or below the midline of the screen (i.e., above or below the pursuit target trajectory). Participants could respond by pressing the up-arrow to indicate that they saw the stimulus above the screens’ midline, or the down-arrow if they perceived the one in the lower half of the screen. To keep the response interval concise, we omitted the final phase of the first experiment in **Experiment 2**, ending after asking if the participants thought they had generated a catch-up saccade (see **Table 1** for an overview of the differences between experiments).

**Table 1.** Stimulus parameters and proportion of stimulus conditions used in Experiments 1–3.

|  |  | Experiment 1 | Experiment 2 | Experiment 3 |
| --- | --- | --- | --- | --- |
| Pursuit target velocity |  | 6, 9, 12 dva/s | 3, 6, 9 dva/s | 3, 6, 9 dva/s |
| Stimulus phase shift velocity |  | 106.65, 109.65, 112.65 dva/s | 56.50, 59.50, 62.50 dva/s | 56.50, 59.50, 62.50 dva/s |
| Stimulus phase shift directions |  | parallel | antiparallel | antiparallel |
| Replayed saccades | First session | Amplitude: 2 dva;<br>Peak vel.: 100.65 dva/s | Amplitude: 1 dva<br>Peak vel.: 53.50 dva/s | Amplitude: 1 dva<br>Peak vel.: 53.50 dva/s |
|  | Later sessions | Amp. 2 dva;<br>Peak vel. 100.65 dva/s | Individualized | Individualized |
| Stimulus conditions<br>(no-stimulus, generated, replayed) |  | 20%, 40%, 40% | 20%, 60%, 20% | 25%, 37.5%, 37.5% |

To examine if participants could be trained to suppress their catch-up saccades (and awareness thereof), participants were instructed in the beginning of the experiment to try and pursue as smoothly as possible. They were additionally informed that perceiving the stimulus likely indicated that a catch-up saccade was (accidentally) generated. Abbreviated instructions were presented before each session as a short reminder.

#### Variations in Experiment 3

The procedure of **Experiment 3** was largely identical to that of **Experiment 2**. We used the same target speeds, stimulus phase shift directions, and response schema (see **Table 1** for an overview of the differences between experiments). All other aspects of the task were kept constant, with the exception of two key changes: (1) the pursuit target was no longer a single dot but a dot cloud composed of 4–8 smaller dots with randomly jittered x- and y-positions to create a horizontally elongated shape, and (2) we introduced trial-wise instruction cues to manipulate the intention behind catch-up saccades. Specifically, participants were asked to either pursue the target as smoothly as possible (unintended saccade condition) or to make an instructed saccade to a briefly highlighted target dot at a defined moment during pursuit (intended saccade condition). These saccade cues appeared either early (200–450 ms after pursuit onset) or late (700–950 ms), allowing us to examine how timing affected compliance with the instruction. The saccade targets were offset horizontally (±0.5, 1, or 1.5 dva) and vertically (±5°) relative to the initial pursuit dot to promote both forward/backward and oblique saccades. This manipulation enabled us to investigate participants’ awareness of both intended and unintended catch-up saccades.

#### Online control of eye positions

During **Experiments 1, 2**, and **3** participants’ eye positions were tracked. Eye and screen coordinates were aligned by conducting standard nine-point calibration and validation procedures before the first trial of each session and whenever necessary. Blinks and deviations in gaze position (>1.5 dva from fixation during the fixation interval, >9 dva from the target dot during the pursuit interval) were likewise monitored in all experiments and led to an abortion of the trial. Aborted trials were repeated at the end of each block in randomized order.

#### Saccade detection

Binocular catch-up saccades were detected in **Experiments 1**, **2**, and **3** using a combination of an acceleration-based threshold and the algorithm described by Engbert and Mergenthaler (2006). For the acceleration-based approach, we sequentially applied low-pass Butterworth filters to the position, velocity, and acceleration data for each component of the binocular eye-tracking signal, using a cutoff frequency of 15 Hz for position and 30 Hz for both velocity and acceleration (c.f., Fooken & Spering, 2020; Harris et al., 2023). If acceleration exceeded a detection threshold of at least 300 dva/s (adjusted upward in cases of lower tracking accuracy) during two consecutive zero-crossing intervals, the corresponding time period was flagged as a potential saccade. If a saccade was simultaneously detected by the velocity-based method within the same interval, the event was classified as a saccade. For velocity-based detection, we used a λ of 5 and a minimum saccade duration of 6 ms (i.e., 3 data samples). To avoid counting fragmented events and reduce false separations, saccadic events were merged if they occurred within 10 ms (i.e., 5 data samples) of one another. Saccade parameters (e.g., saccade onset, amplitude, peak velocity, etc.) were extracted from the velocity-based detection algorithm applied to the raw data, after co-registering the detected events with those identified using the acceleration-based approach.

#### Exclusion of trials from analyses

We excluded saccades (not entire trials) from all analyses if they occurred during the first 500 ms of the fixation period; participants had been instructed to maintain fixation until the target occluded the fixation dot, and only then begin pursuit. Saccade rates were calculated based on this filtered data. For our analyses of visual and saccade sensitivity, we additionally excluded trials in which more than one catch-up saccade was detected. This was done to ensure a reliable estimation of hit and false alarm rates, as the presence of multiple saccades made it unclear whether—and in response to which event—participants provided their response. We also excluded trials in which a replayed catch-up saccade could have rendered the stimulus visible and in which the participant generated at least one (additional) catch-up saccade.

### QUANTIFICATION AND STATISTICAL ANALYSIS

#### Visual sensitivity to intra-saccadic stimulation

##### Eye movement generation

We first estimated observers’ visual sensitivity to the stimulus in all three experiments by examining their responses to the first question asked after each trial: “Did you perceive a STIMULUS FLASH?” Individual hit rates (HIR) were calculated based on affirmative responses in trials in which a stimulus was present. Similarly, individual false alarm rates (FAR) were calculated based on affirmative responses in trials without a stimulus. To assess the effect of eye movement generation and stimulus visibility on visual sensitivity, HIRs were calculated separately for trials with and without generated and replayed catch-up saccades. Due to the low number of false alarms, FARs were calculated separately for generated and replayed eye movements, but irrespective of whether an eye movement was actually generated. In **Experiments 2** and **3**, HIRs and FARs were additionally computed by session (**Exp. 2**) or by eye movement type (intended vs. unintended; **Exp. 3**). All rates were calculated individually for each observer. Sensitivity was then computed by z-transforming individual hit and false alarm rates and subtracting the latter from the former. To assess the effect of target velocity on stimulus visibility, we repeated this analysis separately for each stimulus condition and target velocity (**Exp. 1**: 6, 9, 12 dva/s; **Exps. 2** and **3**: 3, 6, 9 dva/s). This was done separately from the main analysis, as data were insufficient for some participants to robustly estimate sensitivity for each velocity level across all conditions. We predicted that visual sensitivity would be modulated by the presence of an eye movement, with higher sensitivity expected in trials featuring a generated or replayed catch-up saccade compared to those without one. In contrast, we anticipated no sensitivity differences between generated and replayed saccades. Finally, we expected visual sensitivity to be unaffected by target velocity, session (**Exp. 2**) as well as saccade type (**Exp. 3**).

To evaluate whether the stimulus was truly invisible in the absence of a catch-up saccade, we computed average sensitivity indices for trials without (detected saccade) per experiment and examined whether their 95% confidence intervals (CI95%) included zero. We tested whether saccade generation increased visual sensitivity by comparing the CI95% of averaged sensitivities to zero; if the interval excluded zero, the increase was deemed significant. To determine whether visual sensitivity differed between generated and replayed saccades, we conducted a one-way rmANOVA with saccade type (generated vs. replayed) as a within-subject factor (**Exp. 1**), and two-way rmANOVAs that additionally included session (**Exp. 2**) or saccade type (intended vs. unintended; **Exp. 3**) as a second within-subject factor. To assess the effect of target velocity on visual sensitivity, we conducted one-way rmANOVAs with target velocity as a within-subject factor in all experiments.

##### Eye movement kinematics

Because visual sensitivity likely depends on how well the stimulus is stabilized on the retina, we examined sensitivity as a function of retinal velocity during catch-up saccades. Retinal velocity was calculated by subtracting the fixed phase shift speed from each catch-up saccade’s peak velocity, with positive values indicating that eye movement and phase shift directions were identical and negative values indicating that they were oriented in opposite directions. We categorized retinal velocities below 30 dva/s as ‘low’ and those above 30 dva/s as ‘high.’ Hit rates, false alarm rates, and sensitivity measures were computed separately for generated and replayed eye movements, as well as for different. In **Experiment 1**, HIR and

FAR were calculated as in previous analyses: a hit was defined as a “stimulus seen” response in stimulus-present trials, and a false alarm as a “stimulus seen” response in stimulus-absent trials. In **Experiments 2** and **3**, however, the two stimuli were always presented in opposite directions, so we added a second response question during the response phase: participants were asked which stimulus they had seen (i.e., the one above or below the gaze trajectory). Sensitivity measures were based on this second question. Specifically, a hit was defined as a report of the stimulus location for which the retinal velocity was closer to zero, while a false alarm was defined as a report of the opposite stimulus—i.e., the one for which retinal velocity was farther from zero. Visual sensitivity was calculated by subtracting z-transformed FAR from z-transformed HIR as before, but separately for trials with high and low retinal velocities. We predicted that retinal velocity should strongly modulate visual sensitivity, with much higher visual sensitivity in trials with low compared to high retinal velocities.

To assess whether visual sensitivity depended on retinal stabilization (i.e., lower retinal velocities), we conducted separate two-way repeated-measures ANOVAs for each experiment, with retinal velocity (high vs. low) and stimulus condition (generated vs. replayed saccades) as within-subject factors. Due to missing data in at least one condition combination, five participants in **Experiment 2** and two participants in **Experiment 3** were excluded from this analysis.

#### Motor control of catch-up saccades

##### Saccade rates

To assess motor control, we analyzed saccade rates, calculated as the number of saccades divided by the number of trials, and normalized by the average trial duration. Saccade rates were analyzed separately for each experiment and calculated separately based on stimulus presence (present vs. absent), target velocity, as well as session (**Exp. 2**) and saccade type (intended vs. unintended; **Exp. 3**). We predicted that saccade rates would be higher in trials with higher target velocity but would decline across sessions if participants were able to exert conscious motor control over their catch-up saccades. We also expected fewer unintended saccades when participants followed the instruction to pursue, compared to intended catch-up saccades in trials with a saccade instruction. While we did not have specific predictions regarding stimulus presence, we expected (if anything) a stronger training effect—reflected in a greater reduction in saccade rate over sessions—for trials with a stimulus than for those without.

To determine significance, we conducted a two-way rmANOVA with the within-subject factors stimulus presence (present vs. absent) and target velocity for **Experiment 1**. For **Experiments 2** and **3**, we performed separate three-way rmANOVAs that included the additional factor session (**Exp. 2**) or saccade type (intended vs. unintended; **Exp. 3**). To corroborate these results, we conducted equivalent Bayesian model comparisons: These models all models included stimulus presence and target velocity as fixed effects and participant as a random effect. The models for **Experiment 2** and **3** additionally contained factor session (**Exp. 2**) and saccade type (intended vs. unintended; **Exp. 3**). This allowed us to assess the strength of evidence for main effects and interactions beyond traditional significance testing.

One participant had missing data in a single condition combination (highest target velocity in one session) in **Experiment 2**. This participant was excluded from the rmANOVA but retained in the Bayesian model comparison, which accommodates unbalanced data. The Bayesian model converged without issue, using all available data except for the missing cell.

##### Saccade latencies

To examine whether participants showed a more subtle form of training effect beyond changes in saccade rate over time, we additionally calculated the latency of the first saccade in each trial and condition, investigating whether participants were able to delay saccade initiation (i.e., withhold a saccade until later in the trial). Because stimulus presence had no effect on saccade rate and we had no preregistered predictions regarding its influence on latency, we excluded this factor from the analysis. Latencies were therefore calculated separately for target velocity, as well as for session (**Exp. 2**) and saccade type (intended vs. unintended; **Exp. 3**).

To determine statistical significance, we conducted a one-way rmANOVA with target velocity as the sole within-subject factor for **Experiment 1**. For **Experiments 2** and **3**, we performed separate two-way repeated-measures ANOVAs that included target velocity and either session (**Exp. 2**) or saccade type (intended vs. unintended; **Exp. 3**) as within-subject factors. As in the previous analysis, we corroborated these results using equivalent Bayesian model comparisons. All Bayesian models included target velocity as a fixed effect and participant as a random effect. The models for **Experiments 2** and **3** additionally included session (**Exp. 2**) and saccade type (**Exp. 3**) as fixed effects.

As in our analysis of saccade rate, one participant was excluded from the rmANOVA for **Experiment 2** due to missing data but was retained in the Bayesian model comparison, as the model was able to converge.

#### Eye movement sensitivity

Lastly, to assess observers’ awareness of their catch-up saccades, we calculated saccade sensitivity—defined as the ability to judge whether a saccade had been generated in the preceding trial. To this end, we analyzed participants’ responses to the question “Do you think you generated a CATCH-UP SACCADE?” A “yes” response in a trial with an actual catch-up saccade was classified as a hit; the same response in a trial without a saccade was classified as a false alarm. Sensitivity was computed by z-transforming individual hit and false alarm rates and subtracting the latter from the former. This analysis was performed separately for stimulus-present and stimulus-absent trials. Crucially, to control for the influence of (trans-saccadic) visual information—that is, to ensure that saccade detection was not merely driven by visual detection of the stimulus—we adjusted the trial selection for stimulus-present conditions: HIRs were based only on trials in which a generated saccade could have rendered the stimulus visible. FAs, in contrast, were based on replay trials—those without a generated saccade, but in which the replay of a previously generated catch-up saccade could have similarly rendered the stimulus visible. We conducted an additional analysis applying the trial split separately for each target velocity level. We predicted higher saccade sensitivity in stimulus-present compared to stimulus-absent trials, due to the contribution of visual information—particularly in **Experiment 2**, where seeing the stimulus implied a 75% probability that a saccade had been generated—despite our efforts to control for this. In **Experiment 2**, we further expected saccade awareness to increase across sessions if sensorimotor awareness of catch-up saccades could be improved through training. In **Experiment 3**, we predicted above-zero saccade sensitivity for intended saccades and, if intention indeed drives awareness, this sensitivity should exceed that for unintended saccades. We did not pre-register specific predictions for the effect of target velocity.

To assess whether participants were able to detect their own catch-up saccades, we used a one-way rmANOVA with stimulus presence (present vs. absent) as a within-subject factor in **Experiment 1**. For **Experiment 2**, we conducted a two-way rmANOVA with stimulus presence and session as within-subject factors, and similarly, in **Experiment 3**, with stimulus presence and saccade type (intended vs. unintended) as within-subject factors. To complement the frequentist analyses, we performed equivalent Bayesian model comparisons: In **Experiment 1**, the model included stimulus presence as a fixed effect and participant as a random effect. For **Experiment 2**, the model included additionally included the factor session, and for **Experiment 3**, saccade type was included instead of session. To determine the effect of target velocity, we repeated the analyses including target velocity as a factor in the rmANOVA and Bayesian model comparisons—adding it in **Experiment 1,** and replacing session or saccade type with it in **Experiments 2** and **3**.

We excluded one participant from the analysis of the effect of target velocity on saccade sensitivity in **Experiment 1**, and two participants from the equivalent analysis in **Experiment 2**. They were excluded from both the rmANOVA and Bayesian model comparisons because the models failed to converge when their data were included.

## Supplementary Material

### S1: Causal assignment from **Experiment 1**

In **Experiment 1**, at the end of each trial in which observers reported seeing the stimulus, we additionally asked them to provide a certainty rating regarding the causal connection between their eye movement and stimulus perception. Specifically, if observers believed they had generated an eye movement, we asked whether they thought this movement caused the change in stimulus visibility. Conversely, if they believed they had not generated an eye movement, we asked how confident they were that the stimulus was not caused by them. Participants could report on a scale using one of four options: not sure, rather unsure, rather sure, and very sure. We included this question to gain insight into participants’ metacognitive awareness of the relationship between their eye movements and the resulting changes in stimulus visibility. We compared this separately for correct assignments (e.g., when a catch-up saccade was generated, the stimulus was seen, and participants reported making the saccade) and incorrect assignments (e.g., when no eye movement occurred, the stimulus was visible due to a replay, but participants still believed they caused the stimulus perception by generating a saccade).

Participants tended to be rather uncertain about the connection between their eye movements and stimulus visibility: average certainty ratings hovered near zero—the center of the scale and the point of highest uncertainty—regardless of stimulus condition (generated: *mean* = 0.20 ± 0.24; replayed: *mean* = 0.15 ± 0.26) or correctness of the assignment (correct: *mean* = 0.32 ± 0.26; incorrect: *mean* = 0.06 ± 0.24; **Fig. S1**). The two-way repeated-measures ANOVA compared average certainty ratings across assignment correctness (correct vs. incorrect) and stimulus condition (generated vs. replayed). Three participants were excluded from this analysis because reliable certainty ratings could not be calculated across all bins. Results showed that neither factor nor their interaction significantly affected certainty (all *p*s > 0.250, all BF10 < 0.70), indicating that participants’ uncertainty remained consistent across conditions.

Our data indicates that participants’ certainty about the causal connection between their eye movements and stimulus visibility was generally low and unaffected by stimulus type or assignment correctness, suggesting limited metacognitive awareness of the relationship between their actions and perceptual outcomes. Interestingly, participants showed lower certainty and a more heterogeneous response pattern (reflected in larger 95% confidence intervals) when reporting that they had generated an eye movement themselves, regardless of whether this was correct. This again indicates low sensorimotor awareness of saccade generation, suggesting that even when participants believe they caused an eye movement, their confidence in that connection remains weak and variable.

### S2: Observer groups in **Experiment 2**

We were concerned that training participants over just four sessions to suppress their catch-up saccades might be too short for any potential training effects to emerge. To address this concern and incorporate observer experience into our design, we invited two groups to participate in our second experiment: five naïve observers, who had never taken part in an eye-tracking study, and five expert observers from the lab who had participated in several previous experiments. To assess whether eye movement expertise influenced visual sensitivity, motor control, and sensorimotor awareness of saccades, we repeated all major analyses for **Experiment 2** with the additional factor of observer group.

Focusing on visual sensitivity first, we examined how well participants from each group perceived the stimulus in trials without a catch-up saccade and found sensitivity to be close to zero in both groups (naïve: d’ = 0.10 ± 0.43; expert: d’ = 0.05 ± 0.47; **Fig. S2a**). When turning to trials with a saccade, we found saccade sensitivity to be substantially higher in both observer groups (naïve: d’ = 1.34 ± 1.16; expert: d’ = 1.75 ± 0.70; **Fig. S2a**). A two-way mixed-effects ANOVA confirmed a significant overall increase in sensitivity when a saccade occurred (*F* (1,8) = 25.27, *p* = 0.001), with no significant difference between groups (*F* (1,8) = 0.28, *p* > 0.250) and no interaction (*F* (1,8) = 0.77, *p* > 0.250). A second two-way mixed-effects ANOVA showed that the increase in sensitivity was comparable for generated and replayed eye movements (*F* (1,8) = 0.32, *p* > 0.250), and this pattern held across both observer groups (naïve vs. expert: *F* (1,8) = 0.69, *p >* 0.250; interaction: *F* (1,8) = 0.04, *p* > 0.250).

To determine whether experienced participants had greater conscious motor control over their catch-up saccade generation, we first compared average saccade rates between groups. We found similar rates (naïve: 1.52 ± 0.98 s⁻¹; expert: 1.44 ± 0.67 s⁻¹; **Fig. S2b**), with no statistically significant difference between them (*t* (7.0) = -0.19, *p* > 0.250, BF10 = 0.5). To assess potential learning advantages in expert observers, we then calculated saccade rates for each session individually and submitted the data to a two-way mixed-measures ANOVA. This analysis revealed no main effects of session (*F* (3,24) = 0.42, *p* > 0.250) or observer group (*F* (1,8) = 0.03, *p* > 0.250), and no interaction (*F* (3,24) = 1.35, *p* > 0.250), suggesting that saccade suppression performance remained stable over time and did not benefit from prior experience (c.f., **Fig. S2b**). A Bayesian model comparison corroborated these results, with strongest—albeit still low—support for a model including only observer group (BF10 = 0.72), while models including session or interactions were substantially less likely (all BF10 < 0.26).

Lastly, we investigated whether expert observers might be more sensitive to their catch-up saccades or better able to use the saccade-contingent visual feedback to determine whether a saccade had occurred. Irrespective of stimulus presence, saccade sensitivity was similarly low for both naïve (present: d’ = 0.09 ± 0.74; absent: d’ = 0.19 ± 0.73) and expert observers (present: d’ = 0.18 ± 0.81; absent: d’ = 0.03 ± 0.68; **Fig. S2c**). A two-way mixed-measures ANOVA with stimulus presence and observer group as factors confirmed that none of the effects were statistically significant (all *p*s > 0.250). We again corroborated these results using a Bayesian model comparison. The analysis demonstrated that the model including observer group was the best-fitting model (BF10 = 0.56), though it provided only weak evidence relative to the null. Alternative models including stimulus presence (present vs. absent: BF10 = 0.40) or interactions (BF10 < 0.23) showed somewhat to substantially worse fit, indicating that the addition of these factors did not improve model performance. Overall, the evidence for any effect was weak, with Bayes Factors close to 0.5 reflecting only anecdotal support.

Analyses comparing naïve and expert observers revealed no evidence that prior experience with eye-tracking conferred an advantage in visual sensitivity, motor control, or sensorimotor awareness of catch-up saccades. Both groups exhibited similarly low sensitivity and saccade rates, with no evidence of learning effects across sessions. It stands to reason that eye movement expertise does not enhance sensorimotor awareness or control of catch-up saccades. Additionally, neither awareness nor control likely benefits from long-term training or continued exposure to environments with repeated and tightly controlled eye movement behavior (i.e., piloting or participation of psychophysical experiments).

### S3: Saccade parameters in response to target manipulations in **Experiment 3**

An alternative way to investigate how well observers can control saccade generation during pursuit eye movements is to examine how effectively participants adapted their saccades to the instructed eye movements in **Experiment 3**. Participants were instructed to generate saccades over three distances—0.5, 1.0, or 1.5 dva—either in the direction of the moving target or against it, mimicking forward and backward corrective saccades during pursuit. The go-instruction was presented either early (200–450 ms after trial onset) or late (700–950 ms after trial onset). Late go-cues were included to create conditions in which participants were instructed to saccade but had limited time to comply with the instruction. This allowed us to assess saccade rate as a function of instruction timing, target distance, and saccade direction.

We conducted a three-way rmANOVA with saccade direction (forward vs. backward), target distance, and go-cue timing. The analysis revealed a significant main effect of saccade direction (forward vs. backward*: F* (1,9) = 5.56, *p* = 0.043) indicating that participants made slightly but significantly more saccades when instructed to saccade forward (**Fig. S3a**). There was also a significant main effect of target distance (0.5, 1.0, 1.5 dva: *F* (2,18) = 9.10, *p* = 0.002), showing that saccade rate increased with increasing saccade amplitude. The main effect of go-cue timing was significant as well (early vs. late: *F* (1,9) = 16.08, *p* = 0.003), reflecting a higher saccade rate when the instruction was given early compared to late. Finally, we observed a significant interaction between saccade direction and go-cue timing (*F* (1,9) = 5.62, *p* = 0.042), suggesting that the effect of timing differed depending on saccade direction. None of the other interactions were significant (all *p*s > 0.154).

We conducted the same three-way rmANOVA on the first saccade following the go-cue. Here, only two main effects reached significance (**Fig. S3b**): The analysis revealed a significant main effect of saccade direction (forward vs. backward: *F* (1,9) = 11.64, *p* = 0.008), indicating that saccade amplitudes were larger when participants were instructed to saccade forward compared to backward. There was also a significant main effect of target distance (0.5, 1.0, 1.5 dva: *F* (2,18) = 13.80, *p <* 0.001), showing that saccade amplitude increased in line with the instructed target distance. Unlike in our previous ANOVA for saccade rate, the timing of the saccade go-cue had no effect (*F* (1,9) = 0.66, *p* > 0.250), nor were there any significant interactions (all p-values > 0.250).

Together, these analyses show that while saccade rate is influenced by the timing of the go-cue, saccade direction, and target distance, saccade amplitude is primarily driven by saccade direction and target distance but unaffected by go-cue timing. These results provide further evidence that saccade control is possible but primarily shaped by low-level visual factors such as target distance and direction.

### S4: A closer look at saccade sensitivity: Hit and false alarm rates across experiments

To gain a more nuanced understanding of saccade awareness, we analyzed hit rates (saccade detection) and false alarm rates (erroneous reports in the absence of a saccade) separately. While a sensitivity measure is better suited to capturing true sensorimotor awareness of catch-up saccades, it does not reveal whether effects were driven by detection accuracy, guessing, or changes in participants’ caution. It might also fail to capture potential effects of the additional factors we investigated—stimulus presence, training, and intention.

To investigate this, we conducted a two-way rmANOVA with the type of response rate and stimulus as factors for **Experiment 1**. The analysis revealed a significant main effect of stimulus presence (present vs absent: *F* (1,7) = 9.02, *p* = 0.020), indicating rates were higher when the stimulus was present (hit = 0.18 ± 0.13; fa = 0.15 ± 0.09) compared to stimulus-absent trials (hit = 0.11 ± 0.10; fa = 0.09 ± 0.08). There was no main effect of rate type (hit vs. fa: *F* (1,7) = 0.59, *p* > 0.250), suggesting that overall false alarm rates were not significantly different from hit rates. We also found no significant interaction (*F* (1,7) = 0.60, *p* > 0.250), indicating that the effect of stimulus presence was similar across both types of responses (**Fig. S4**). These results are broadly supported by our Bayesian model comparison: there was moderate evidence for a model including stimulus presence (BF10 = 5.45), anecdotal evidence against a main effect of response rate type (BF10 = 0.46), and no clear evidence for an interaction (BF10 = 1.13).

To examine what affected response rates in **Experiment 2**, we conducted a three-way rmANOVA with the same two factors as before (stimulus presence and rate type), and session number (to assess training effects) as factors. We found that the main effect of session approached significance (*F* (2.2,19.4) = 3.34, *p* = 0.053 after Greenhouse-Geisser correction for violation of sphericity), as did the main effect of condition (*F* (1,9) = 4.81, *p* = 0.056) and their interaction (*F* (1.9,17.4) = 3.07, *p* = 0.074 after Greenhouse-Geisser correction for violation of sphericity). This pattern is consistent with that observed in the first experiment: in **Experiment 2**, we again found higher response rates during stimulus-present trials (hit = 0.23 ± 0.15; false alarm = 0.20 ± 0.13) than during stimulus-absent trials (hit = 0.09 ± 0.06; false alarm = 0.11 ± 0.07), although these differences did not reach statistical significance. Moreover, hit and false alarm rates did not significantly differ from one another (*F* (1,9) = 0.03, *p* > 0.250), suggesting that stimulus presence influenced overall response tendency rather than selectively affecting detection or guessing (see **Fig. S4**). All remaining interactions remained insignificant (all *p*s > 0.069 after Greenhouse-Geisser correction for violation of sphericity). To complement these effects, we conducted a Bayesian model comparison. The model including **session** and participant received the strongest support (BF10 = 1.71 × 10^31^), with similar evidence for models that additionally included stimulus presence (BF10 = 1.06 × 10^31^) or response type (BF10 = 4.67 × 10^30^). More complex models with interaction terms performed substantially worse, and models omitting **session** entirely yielded very low Bayes factors (all BF10 < 0.26), indicating that these factors alone poorly accounted for the observed data.

To explore which factors influenced response rates in **Experiment 3**, we conducted a three-way rmANOVA with the same two factors as before (stimulus presence and rate type), and this time included intention (manipulated via pursuit and saccade instruction) to assess the role of volition. The analysis revealed a robust main effect of intention (*F* (1,9) = 88.16, *p* < 0.001), indicating that response rates differed markedly depending on whether the eye movement was intentional (hit = 0.82 ± 0.18; FA = 0.80 ± 0.18) or unintentional (hit = 0.11 ± 0.08; FA = 0.08 ± 0.06). The main effect of stimulus presence yet again trended but failed to reach significance (*F* (1,9) = 4.31, *p* = 0.068), resembling the data of the first two experiments: participants responded more often on stimulus-present trials (hit = 0.49 ± 0.13; FA = 0.46 ± 0.11) than on stimulus-absent ones (hit = 0.43 ± 0.11; FA = 0.42 ± 0.10). Again, the absence of a main effect of rate type (*F* (1,9) = 1.54, *p* > 0.250) suggests that this effect applied similarly to both hits and false alarms, pointing to a general modulation of response likelihood rather than selective changes in detection or guessing. None of the interactions reached significance (all *ps* > 0.250; c.f. **Fig. S4**). We again conducted a Bayesian model comparison to corroborate the results of the rmANOVA. Our analysis revealed strongest support for models including saccade type and participant (BF10 = 1.7 × 10^31^), with models additionally including condition also receiving substantial support (BF10 = 1.1 × 10^31^). Models including only stimulus presence (BF10 = 0.26) or response type (BF10 = 0.24) showed considerably less support, indicating that saccade type was the primary factor influencing response rates.

Across all three experiments, response rates were consistently influenced by task-related factors rather than by differences in detection. In **Experiment 1**, we found higher response rates when the stimulus was present, regardless of whether the response was a hit or a false alarm, suggesting that visibility alone increased participants’ tendency to report a saccade. **Experiment 2** produced a similar stimulus-driven pattern that varied over time. **Experiment 3**, in turn, revealed a strong effect of intention: participants responded far more often when eye movements were instructed and thus intentional, again with comparable rates for hits and false alarms. Crucially, across all analyses, we found no significant main effects of rate type, indicating that the factors manipulated in the task modulated overall response likelihood rather than selectively affecting perceptual sensitivity. Our data, hence, suggests that awareness of catch-up saccades largely reflect expectations shaped by context and intention, rather than precise introspective access to individual eye movements.

---

**Fig M1.**
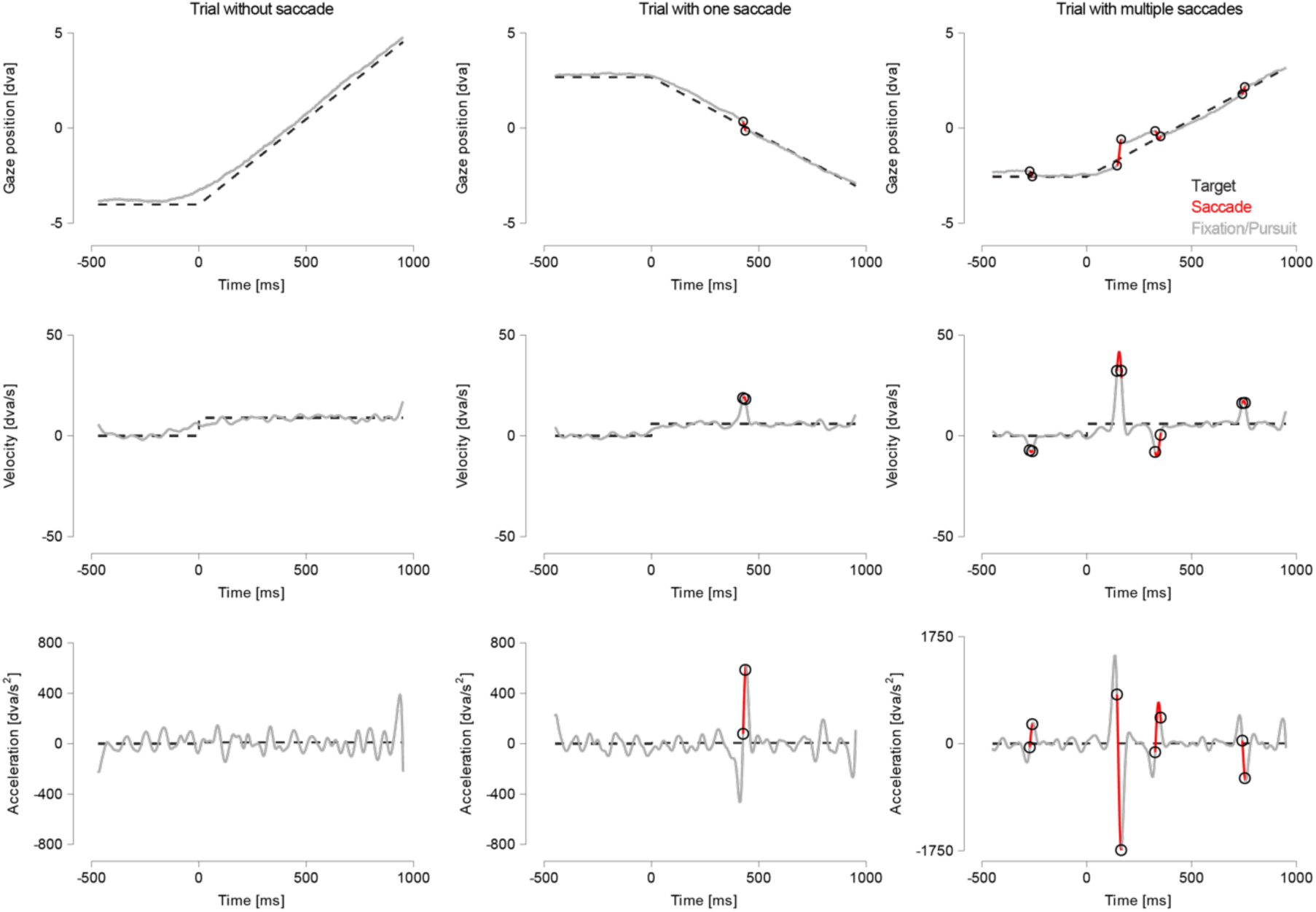
Detection of zero (left), one (middle), or multiple saccades (right column) by the co-registered detection approach. Data show raw position (top), velocity derived from smoothed positions (middle), and acceleration computed from these velocities before final smoothing (bottom) of three randomly selected trials of one participant. Note that the y-axis of the rightmost acceleration plot differs from the others due to the large magnitude of the negative deflection.

**Fig S1.**
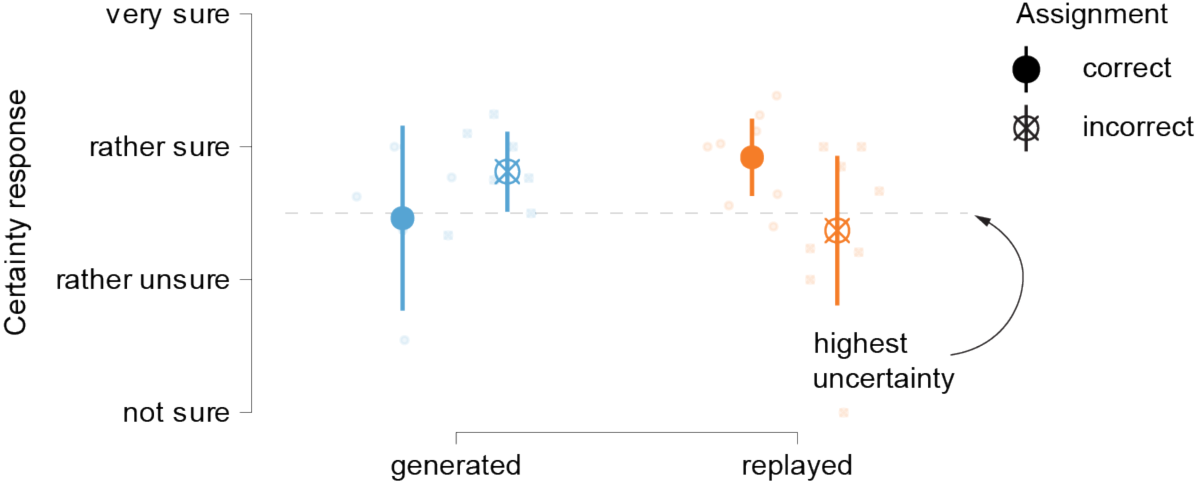
Low certainty about the causal connection between their eye movements and stimulus visibility—for generated and replayed saccades and irrespective of assignment correctness. Error bars represent 95% confidence intervals.

**Fig S2.**
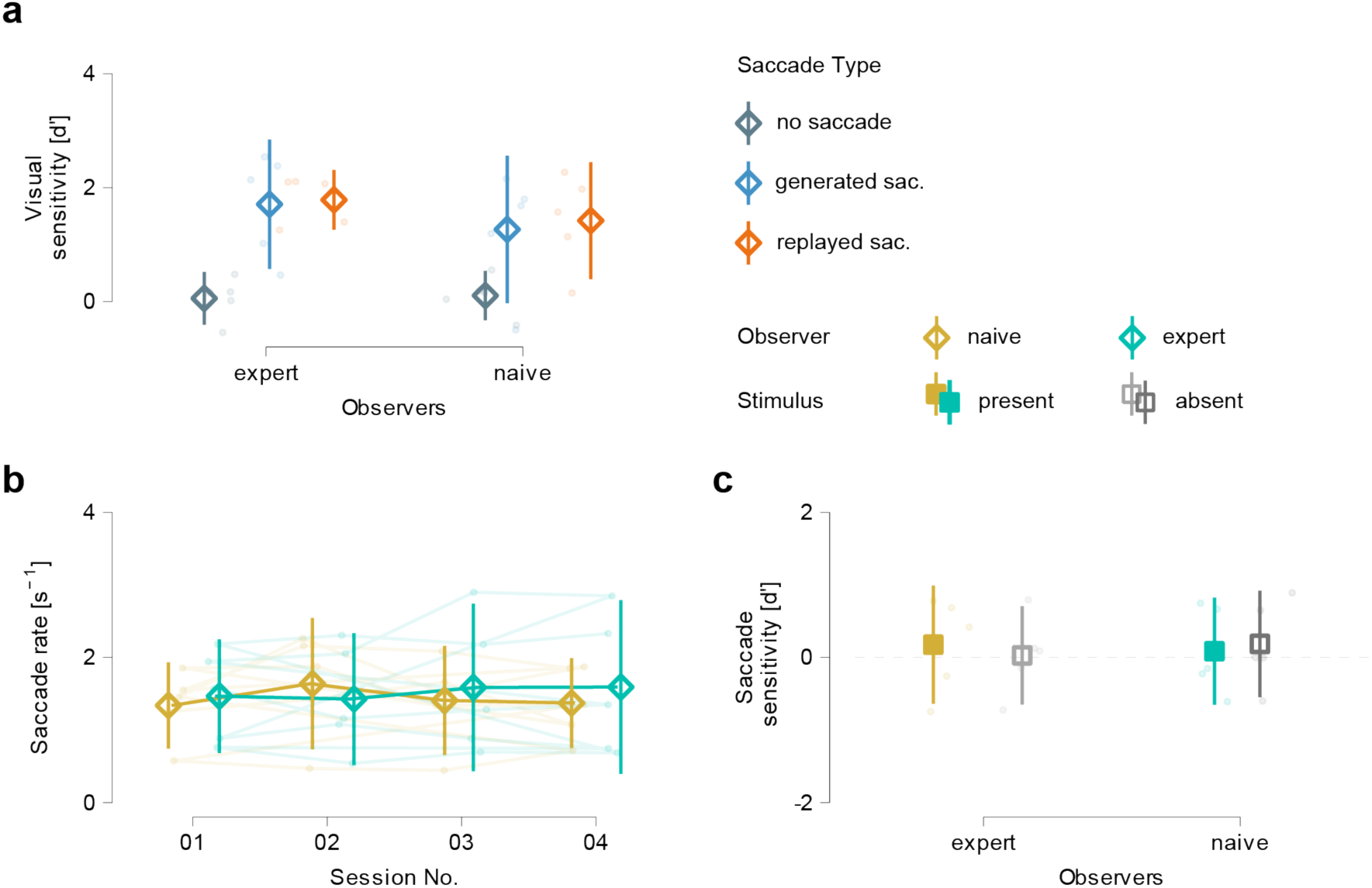
Pre-training level of the observer has no effect on visual sensitivity, motor control, or saccade sensitivity. **a** Visual sensitivity to the stimulus as a function of saccade generation and eye movement condition. Data are shown separately for participants with different pre-training levels: naïve and expert. **b** Development of saccade rate (as an index of motor control training) across the four experimental sessions and separately for naïve and expert observers. **c** Saccade sensitivity as a function of stimulus presence and pre-training level. All panels: Error bars represent 95% confidence intervals.

**Fig. S3.**
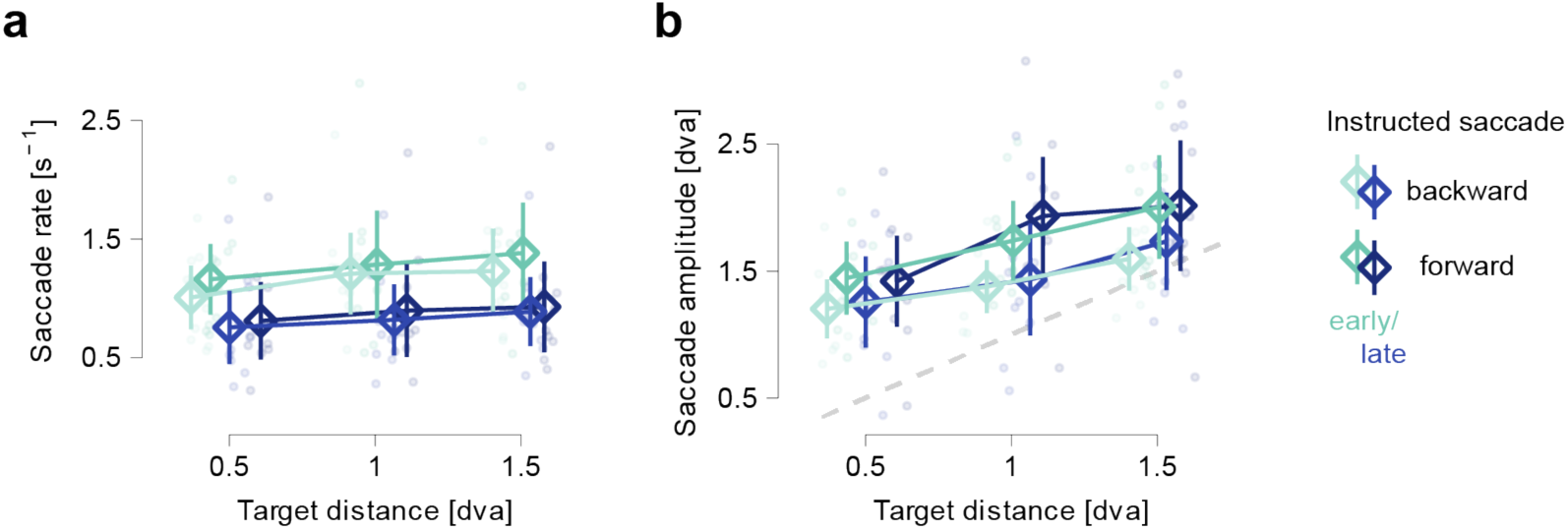
Saccade rate and amplitude increase with target distance in instructed catch-up saccade trials. **a** Saccade rate as a function of the timing of the go-instruction (early vs. late), the instructed target distance (0.5, 1.0, 1.5 dva), and saccade direction (forward vs. backward) in **Experiment 3**. **b** Saccade amplitude as a function of the same factors. Dashed line marks the origin (x=y). Both panels: Error bars represent 95% confidence intervals.

**Fig. S4.**
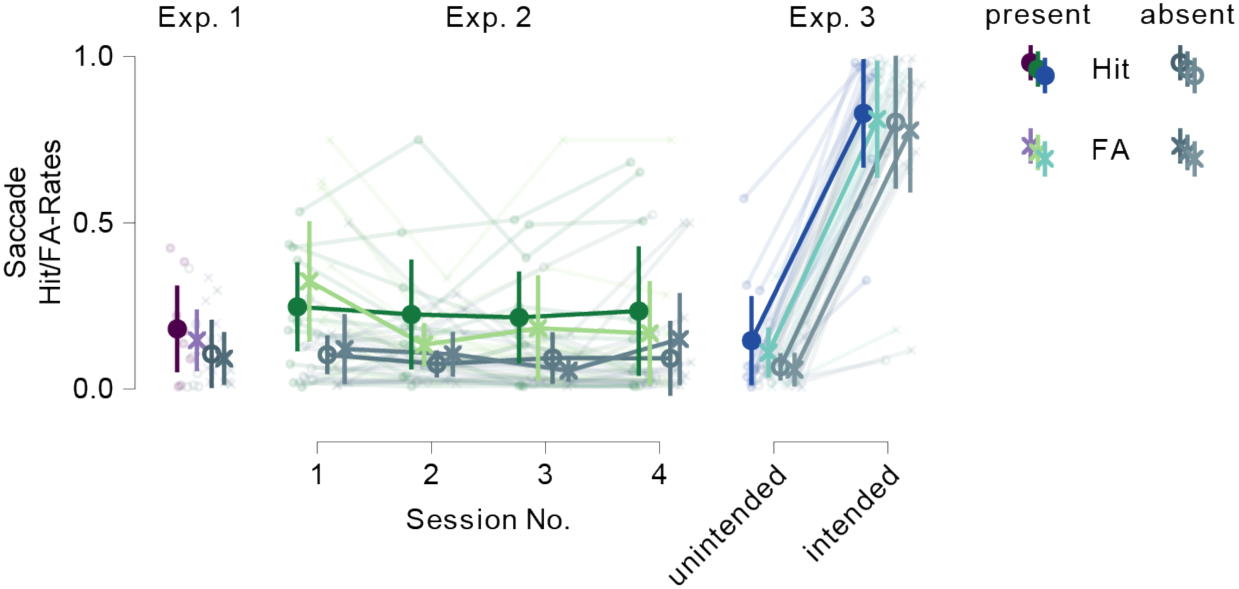
Response rates (hit and false alarms) are similarly affected by stimulus presence and intention. Hit and false alarm rates as a function of stimulus presence, how it develops over time to assess training (**Exp. 2**) and is affected by intention (**Exp. 3**).

## Footnotes

* Assuming an average coasting velocity of 60–85 mm s−1 (c.f., Wu et al., 2007) and an observer distance between 2.5 and 10 meters, koi fish travel at approximately 0.3–2 dva/s.

